# Fitness effects of new mutations are small and heavily confounded with non-genetic sources of variation in Escherichia coli

**DOI:** 10.64898/2026.08.23.746582

**Authors:** Julie M. Grosse-Sommer, Dhobasheni Newman, Jarrod D. Hadfield

**Affiliations:** Institute of Ecology and Evolution, School of Biological Sciences, University of Edinburgh, Edinburgh, UK

**Keywords:** Mutation, distribution of fitness effects, microbial evolution, environmental effects

## Abstract

The distribution of fitness effects (DFE) of new mutations underpins our understanding of molecular evolution, mutation load and the maintenance of quantitative genetic variation. Most direct estimates in microbes rely on mutation-accumulation (MA) lines that harbour many mutations, so that only the mean and variance of the DFE can be inferred reliably and non-genetic (‘environmental’) differences between lines are usually ignored. Here we generated 192 Escherichia coli MA lines that accumulated no mutations, one mutation or more than one mutation in similar proportions. Comparing the growth rates of lines with mutations with the mutation-free controls allowed us to partition fitness variation into genetic and non-genetic components. We estimate the average selection coefficient of a single mutation is unlikely to be less than −0.0020 (with 95% credibility) - a value that is small, not significantly different from zero, and largely concordant with most previous estimates from multi-mutant MA studies, although those earlier results have often been misinterpreted as implying much larger deleterious effects. Crucially, systematic non-genetic effects among lines were an order of magnitude larger than mutational effects, and failure to model them would have biased the mean selection coefficient downwards by an order of magnitude. Our study demonstrates that spontaneous mutations in E. coli are typically only mildly deleterious and that rigorous controls for environmental effects are essential for unbiased inference of the DFE in microbial systems.

## Introduction

The distribution of fitness effects (DFE) characterizes the probability that a new mutation has a given fitness effect. The DFE plays a central role in population and quantitative genetics and is informative about the pace at which Muller’s ratchet increases the deleterious load of a population (Haigh, 1978), the maintenance of heritable quantitative genetic variation (Walsh and Lynch, 2018), the rate of molecular evolution (Lanfear et al., 2014), the evolution of sex and recombination (Peck et al., 1997) and the chance that populations with small effective population size survive (Lande, 1994). There is a general consensus that the frequency of deleterious mutations outweighs that of beneficial mutations, and the majority of mutations are of small effect, particularly in organisms with large amounts of non-coding DNA. However, more detailed descriptions of the DFE, such as the average fitness effect of deleterious mutations, the proportion of mutations that are deleterious, neutral or beneficial, or the shape of the distribution of deleterious mutations, are more poorly understood. The large range of proposed distributional models for the DFE bears testimony to this (Eyre-Walker and Keightley, 2007).

Inferring the DFE can be done using sequence polymorphism and divergence data or by directly measuring the fitness of newly introduced mutations. Inference methods using polymorphism and divergence data provide little information about the tails of the DFE, as strongly selected mutations are expected to segregate at a range of extreme frequencies that are hard to distinguish from each other. Even for mutations with more modest fitness effects, inferences about their distribution often make a set of assumptions that may be unrealistic, for example by assuming beneficial mutations do not contribute to polymorphism or neglecting demographic change (Tataru et al., 2017; Moutinho et al., 2019). While the latter assumption can be relaxed by the inclusion of a comparative set of neutral sites, new assumptions have to be made about the neutrality of these sites. For example, it is common to use synonymous sites as a set of neutral sites (Moutinho et al., 2019), although it is well known that these sites can be under selection (Rahman et al., 2021; Bailey et al., 2021).

To avoid the restrictive assumptions detailed above, and to measure the frequency of larger fitness effects more precisely, direct inferences of the DFE using mutation accumulation (MA) lines can be useful. By propagating lines at very low *N*_*e*_, ideally *N*_*e*_ = 1, the fixation of new mutations is dominated by stochasticity rather than by their fitness effect and a representative sample of mutations can be sampled (Halligan and Keightley, 2009). As it is difficult to make precise inferences on the properties of the DFE from few mutations, mutation rates are often artificially elevated when working with organisms with low genomic mutation rates, such as microbes, either by using external mutagens (Elena et al., 1998; Sanjuán et al., 2004; Wloch et al., 2001) or by using lines that have deficient DNA mismatch repair mechanisms (Wloch et al., 2001). However, the resulting spectrum of mutations (Foster et al., 2015) and their corresponding fitness effects (Sane et al., 2023; Wloch et al., 2001) may not be representative of naturally arising mutations. For example, mutations caused by transposable elements often lead to a complete loss of function when inserted into a gene (Couce et al., 2024) and are therefore expected to have larger fitness effects than naturally occurring spontaneous mutations that tend to be base-pair substitutions. In most studies, a modest number of MA lines are used where each line accumulates many mutations, either naturally or through an elevated mutation rate (Ness et al., 2015; Huang et al., 2016; Tenaillon et al., 2016). Under this design, only the aggregate effect of all accumulated mutations in a line can be measured, and so under the central limit theorem we expect the distribution of line effects to become (log) normal as the number of mutations increases. Consequently, as the number of mutations per line increases, the mean and variance are generally the only aspects of the DFE that can be inferred precisely (Keightley, 1994; Mukai, 1964), and even then, accurate estimates rely on there being no directional epistasis. Generating recombinant lines between the ancestor and MA lines has been used to leverage more information about the effects of individual mutations rather than their aggregate effects (Böndel et al., 2019; Crombie et al., 2024). However, to substantially increase the amount of information when multiple mutations occur per line, a large number of recombinant lines is likely required. For organisms with a low genomic mutation rate, an obvious alternative is to run many MA lines, each accumulating only a few mutations, ideally one. To the best of our knowledge, only one study has measured the fitness effects of single spontaneous mutations - Sane et al. (2018) measured the growth rate of single-mutant *E. coli* lines in eleven different carbon environments (see Sane et al. (2023) also).

A final concern with the MA approach is the possible presence of systematic variation in fitness between lines that is not attributable to the newly accumulated mutations, which, if not accounted for, will bias any inference of the DFE. In non-clonal diploid organisms, any residual heterozygosity contributes to the between-line variance as these remaining polymorphisms fix in different lines. In such cases, inferences can be made from the *changes* in the distribution of line effects that occur during MA, although only later MA generations can be used if the ancestor is not sufficiently inbred and the initial residual heterozygosity exceeds the equilibrium within-line heterozygosity (Mackay et al., 1992). In addition, between-line variation can also be caused by persistent differences in the environments of different MA lines. While this is well acknowledged in work on multicellular organisms, where experimental designs and statistical approaches have been employed to minimise or control for it (Shaw et al., 2000; Lynch, 1985), microbial MA studies have traditionally assumed any environmental between-line variation to be negligible. However, transgenerational ‘maternal’ effects are expected to be strong in organisms that primarily reproduce through binary fission (Robert et al., 2018) and if not accounted for are likely to be mistaken for the effects of mutations accumulated in the MA lines (Lynch, 1985).

By propagating 220 *E. coli* MA lines for 36 days, the average time it takes for a single mutation to accumulate, we produced a set of lines that differ by one spontaneous mutation in order to estimate the DFE. The protocol also produced a set of genetically identical lines that did not accumulate any mutations (‘zero-mutant MA lines’), from which the distribution of non-genetic between-line effects could be estimated and controlled for. We find that the mean selection coefficient is small in magnitude, consistent with previous estimates from multi-mutant lines. However, systematic non-genetic effects on fitness were substantially larger, and had we not been able to correct for them, our estimates of the DFE would have been severely biased. Our study highlights the importance of including proper controls in order to differentiate between the effects of new mutations and confounding non-genetic effects on fitness.

## Methods

### Mutation accumulation procedure

*Escherichia coli* K-12 MG1655 strains were originally obtained from the E. coli Genetic Stock Center and kindly provided by Meriem El Karoui. This ‘ancestral strain’ was grown for 4 hours in Miller’s Luria Broth (LB), before being frozen at an equal volume of 50% glycerol at −70°C. Mutation accumulation lines were started by streaking out 220 new lines from the ancestor onto Miller LB Agar. Plates were subsequently put into a stationary incubator at 37° C. After 24h, lines were subject to bottlenecks by picking a single colony from a previously determined spot on the plate, to ensure unbiased sampling, and further propagated on new Miller LB Agar media, following the same procedure as outlined above. In the event that a line did not produce a single isolated colony or did not grow at all, lines used from the previous day that had been stored at 4°C were used for re-streaking. Every two weeks, single colonies were frozen in 50% glycerol and stored at −80°C.

Lines were propagated for 36 days - the length of time we predicted one mutation per line to accumulate, assuming 28 generations per day (Lee et al., 2012) and an estimated mutation rate of 1.0 *×* 10^−3^ per genome per generation for the MG1655-derivative *E. coli* strain PFM2 on LB media (Lee et al., 2012). Assuming the number of mutations per line is Poisson distributed (Lee et al., 2012; Robert et al., 2018), we expected that 37% of strains would acquire one mutation by the end of the experiment. Human error (six lines), extinction (one line) and potential contamination (one line) caused eight out of the 220 lines to not complete the full mutation accumulation cycle.

### Sequencing, read alignment and mutation calling

In addition to the ancestral line, which was sequenced twice, 194 out of the 212 remaining MA lines were selected for sequencing, determined by the quality of their DNA extractions. DNA extractions were performed with Qiagen DNeasy Blood and Tissue Extraction Kit (96-wells method) and stored at −20^*◦*^C. DNA was sequenced at the Centre for Genomic Research, University of Liverpool, with NovaSeq 6000 (2×150), generating an average coverage of 97.1X per line (range 69.0–133.5). Read integrity and quality was assessed with FastQC and MultiFastQC, where all criteria were passed. The raw sequence data for all lines has been uploaded to the NCBI Sequence Read Archive under BioProject ID PRJNA1466858. The following pipeline is described in detail in Mutation Calling SI.

Reads were aligned to the MG1655-K12 E. coli genome (NCBI RefSeq accession NC 000913.3, assembly GCF 000005845.2 (ASM584v2)) using the bwa-mem package (v0.7.19) (Li, 2013) with default parameters. PCR duplicates, secondary reads and non-mapped reads were excluded using samtools (v1.3.1) (Danecek et al., 2021). Base-pair substitutions (bps) and short insertion/deletions (INDELs) were called with bcftools (v1.23.1) (Danecek et al., 2021). Default parameters were used except extended BAQ was used to reduce false-positive INDEL calls and reads with mapping quality *<* 30 and bases with base-quality *<* 20 were excluded. Variants at sites with a Phred-scaled score (QUAL) *<* 20 were excluded leaving 643 variants. All lines, including the ancestor, had two mutations compared to the reference genome: a base change from G to A at position 4,381,253 in the gene *frdA*, a fumarate reductase flavoprotein subunit, and a CG insertion at position 4,296,380 in a non-promoter non-coding region. Excluding these left us with 255 possible mutations, of which 194 had *≥* 30 and/or *>* 95% reads supporting the alternate allele and were considered real. The remaining 61 ‘provisional mutations’ were visualised in the Integrative Genomics Viewer (IGV) (Robinson et al., 2011) and 30 were considered real. Eleven of the provisional mutations were in rRNA genes or in insertion sequence, and 9 were in the *icd* gene of a single line. All but one was considered to be due to mapping error. Of the remaining 41 provisional mutations, 11 were bps and 30 were INDELS. Three of the bps were excluded. Two had less than three reads in support of the mutation and were removed. One had 67 reads in support of the mutation, but 13 reads in support of the reference. We tentatively excluded this mutation, although it could be real and the sequencing sample had some contamination. Nine of the INDELS were also excluded. Seven of those were found in two regions across multiple lines. In the remaining two cases, the provisional mutation only had 2 reads in support of the alternate or only had support in 22% of reads. Of the 21 provisional INDELS considered real, all but 5 fell in a repetitive region where reads supporting the reference did not span sufficient repeats to call the mutation. The remaining 5 INDELS occurred in a single line that had 17 mutations passing the initial filter, one of which occurred in the *mutS* gene (a P to Q amino acid change caused by a mutation at position 2,858,801) which encodes for a DNA mismatch repair protein and is essential for DNA repair (Li, 2007). Given the provisional mutations in this line had several reads in support of the reference, without evidence of contamination at other mutated sites, it was concluded that these mutations occurred after mutation accumulation, during the growth stage prior to sequencing. This line was excluded from further analyses. A single mutation that passed the strict filter was also excluded as it fell in the *icd* gene in the same line in which many such mutations were found (see below). Additionally, two lines had the same single mutation, likely due to a contamination event (they were propagated on the same plate), and one of these lines was removed from further analysis.

Large structural variants (LSVs), namely larger deletions, insertions, inversions or translocations, were called using Parliament2 (Zarate et al., 2021), where LSVs were called by compiling the results of the programs Lumpy, Breakdancer and CNVNATOR and were also manually checked in the Integrative Genomics Viewer (IGV) (Robinson et al., 2011). Three large deletions were found: 321bp (1386588-1386909), 15kb (1196300-1211412) and 456bp (2817640-

2818096). The large 15kb deletion spans *icd* in the sample where many mapping errors appeared in the list of provisional mutations close to this gene. The deletion coincides with the cryptic prophage e14 which has previously been shown to excise on induction of the SOS pathway (Brody et al., 1985). Mutations were annotated using the Bioconductor (Huber et al., 2015) libraries VariantAnnotation (v1.56) (Obenchain et al., 2014), GenomicFeatures (v1.62) (Lawrence et al., 2013) and rtracklayer (v1.70.1) (Lawrence et al., 2009) for R (v4.5.1) (R Core Team, 2025). GPT-5.5 was used when developing the bioinformatic pipeline.

A total of 203 unique mutations were found to have accumulated over the 192 non-excluded lines (see Table 1) with 176 bps (86.7% of all mutations), 24 INDELs (11.8%) and 3 LSVs (1.5%). 133 mutations (75.6% of bps) were coding (excluding pseudogenes) and of these, 101 (75.9%) were non-synonymous. A total of 65 lines (33.9% of lines) accumulated one mutation, while 55 (28.6%) accumulated more than one mutation. 72 lines (37.5%) did not accumulate any mutations. By fitting a quasipoisson generalised linear model to the number of mutations per line we obtained a total per genome per generation mutation rate estimate of 10.489[9.041 – 12.084]*×*10^−4^ using the generation time estimated by Lee et al. (2012). For bps the mutation rate was 9.094[7.743 – 10.593]*×*10^−4^. Both estimates are comparable to those reported by Lee et al. (2012) (1.014*×*10^−3^ and 0.930*×*10^−3^, respectively). The mutation counts across non-excluded lines had a variance/mean ratio of 1.111, close to the Poisson expectation of one. In order to assess the false negative rate of our mutation calling pipeline, we mapped the reads from our 196 libraries to modified reference genomes. For each library three modifications were made to the reference genome: a base was changed, a single base was added and a single base was removed. Our variant calling pipeline was then used to call variants with respect to the modified reference genomes, where the three changes should appear as base-pair substitutions, deletions and insertions, respectively. 100% of bps, 99% of deletions and 98% of insertions were called. As was seen when calling actual mutations, the number of calls that passed our strict filter was lower for deletions (80.4%) and insertions (74%) than for bps (99%).

**Table 1.** Number of lines with count of accumulated mutations.

|  | 0 | 1 | 2 | 3 | 4 | 5 |
| --- | --- | --- | --- | --- | --- | --- |
| Number of Lines | 72 | 65 | 32 | 20 | 1 | 2 |

### Fitness Measurements

Growth rate was measured for 120 non-excluded lines that accumulated at least one mutation, the hypermutator line (excluded from downstream analyses), the ancestral strain, and 68 (out of 72) randomly selected lines that had accumulated no mutations. The 188 MA lines and four ancestral replicate lines were split into two sets, each containing 94 MA lines and 2 replicates of the ancestral line. For each set, the lines were defrosted and grown while shaking for 16 hours at 37^*◦*^C in 10ml LB media, after which 10*μ*L of culture was transferred to a 96-well filled with 190*μ*L of LB. The plate was subsequently shaken for four hours at 37^*◦*^*C*, with optical density (OD) readings taken every five minutes. Over four weeks, a total of 14 replicates were measured, where each replicate was taken from an independent thawing. The membership of each set and the position of each line on the plate was randomised between replicates to ensure well effects were not confounded with line effects. The difference in log OD between consecutive time points was averaged over a sliding window of three time points, and fitness was taken to be the maximum of these averaged differences using gcplyr (Blazanin, 2024) in R (R Core Team, 2025).

Thirty one replicates that were localised on two plates were discarded because they had very high OD measurements, likely due to plate manufacturing error or OD machine malfunction. Two replicates with high OD were discarded because they were inoculated twice. Due to an error of judgement, 40 replicates which had extremely low growth rates prior to plate inoculation were not measured. Certain lines were overrepresented among these replicates, but there was no indication that those lines had unusually low growth rates (measured from their replicates for which growth rate was measured) or that they were caused by mutations in those lines. In addition, one replicate was removed because no growth was detected and one replicate was removed because the estimated growth rate was an outlier due to an anomalous OD reading (see Fitness Analyses SI for details of the filtering).

### Analysis

Our original intention was to model the distribution of mutation effects using a multimembership model and a skew-t distribution (Azzalini and Capitanio, 2003; Pick et al., 2022) in order to allow flexibility in the estimated shape of the DFE. However, prior to this, we fitted simpler linear mixed models to the data that assumed a Gaussian distribution for the line effects and found little evidence that the DFE differed from a point mass at zero. Since more flexible DFE models would likely not be identifiable, we simply report the results from this model.

The fitness measurement (maximum growth rate measured per hour) for replicate *j* in line *i* (*r*_*ij*_) was analyzed using a linear mixed effect model fitted using lmer (Bates et al., 2015). The part of the model relating to biological processes of interest is:

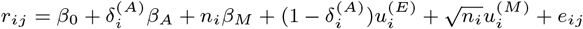

The fixed effect *β*_0_ measures the growth rate of MA lines that have not accumulated any mutations. We refer to this value as the reference growth rate, as all other effects are in respect to this value. While the ancestor’s growth rate could be used as a reference value, we argue that the greater replication of the zero-mutant MA lines provides a more precise estimate than that of the single ancestral strain. 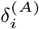 is an indicator variable specifying whether line *i* is the ancestor (*δ*^(*A*)^ = 1) or an MA line (*δ*^(*A*)^ = 0) such that *β*_*A*_ measures how much the fitness of the ancestor differs from the reference or zero-mutant MA lines, on average. *n*_*i*_ is the number of mutations MA line *i* has accumulated, and so *β*_*M*_ measures the average fitness effect of each mutation assuming a linear relationship with the number of mutations (i.e. no directional epistasis). *u*^(*E*)^ are line-specific random effects (excluding the ancestral line) with zero mean and estimated variance 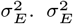 captures the systematic between-line variance in fitness that is not due to accumulated mutations. This variance can be estimated from the 68 lines that did not accumulate any mutations in the experimental process, since they are genetically identical, and any consistent difference in fitness between these lines must be due to non-genetic (environmental) effects. 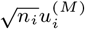 is also a line-specific effect (for line *i*) that can be attributed to mutations. *u*^(*M*)^ are fitted as random with zero mean and estimated variance 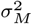. The marginal distribution of 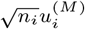therefore has variance 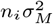 such that the between-line variance attributable to this term increases linearly with the number of mutations, and is absent for the ancestor and MA lines that accumulated no mutations. The between-line variance due to mutations is expected to increase linearly with the number of mutations in the absence of directional epistasis. The total between-line variance in fitness for an MA line with *n* mutations is therefore 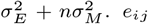 is the residual for the *j*^*th*^ replicate of line *i* and is assumed normal with zero mean and variance 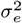 : the within-line variance due to environmental effects and measurement error. In addition, terms were included to account for technical variation. Differences in growth between the inner and outer wells, and between assays inoculated by different people, were fitted as fixed, whereas plate, plate by column and plate by row effects were fitted as random (see Fitness Analyses SI for code).

The significance of the fixed effects was assessed using t-tests with the degrees of freedom obtained using Satterthwaite’s (1946) method, as implemented in lmerTest (Kuznetsova et al., 2017). The significance of the random effect variances were assessed using likelihood ratio tests. Confidence intervals were obtained using profile likelihood.

### Relative Fitness and Selection Coefficients

Under density independence, our fitness measurements are equivalent to the Malthusian parameter (Chevin, 2011). Using the notation *r*_*w*_ to indicate the Malthusian parameter of the reference ‘wild-type’ and *r*_*i*_ to indicate the Malthusian parameter of MA line *i* due to the accumulated mutations, we have the equations:

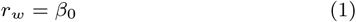

and

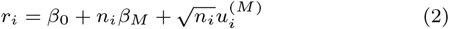

The selection coefficient for MA line *i* is *r*_*i*_ *− r*_*w*_, although this is in units *per hour* and is therefore hard to compare across studies and species (Chevin’s Equation 2.2, Fink and Manhart’s Equation 7). To make selection coefficients more comparable, they can be put on the scale of generations (Chevin (2011), but see Fink and Manhart (2024)). *E. coli* reproduces by binary fission, so assuming no cell death, which is believed to be negligible (Stewart et al., 2005), the generation time of the reference can be expressed as *ln*(2)*/r*_*w*_ (Chevin, 2011). Consequently, the selection coefficient for line *i* scaled by generation time can be obtained by using Equation 3.2 in Chevin (2011):

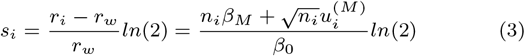

When referring to selection coefficients in what follows, we are referring to these generation-scaled selection coefficients, where the average generation time of the zero-mutant lines is estimated to be *ln*(2)*/β*_0_ = 38 minutes. Since the line effects, *u*^(*M*)^, are assumed normal the selection coefficients for single mutations are also assumed normal with mean

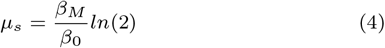

and variance

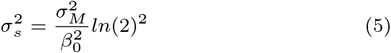

Since the mean and variance are all non-linear functions of model parameters, we obtained credible intervals on these derived parameters by refitting the model described above using MCMCglmm (Hadfield, 2010) with *F*_1,1_ priors scaled by 10 on the random effect variances and a flat prior on the residual. Posterior modes from this model aligned closely with the estimates obtained from lmer. In addition, we also calculated the mean selection coefficient had the ancestor been treated as the reference (i.e *r*_*w*_ = *β*_0_ + *β*_*A*_): *ln*(2)(*β*_*M*_ *− β*_*A*_)*/*(*β*_0_ + *β*_*A*_).

### Comparison with other studies

To compare our results with a similar dataset, we utilized data from lines containing single spontaneous mutations in *E. coli* grown in LB (Sane et al., 2018). The mutation accumulation protocol mirrored ours, with the primary difference being that fewer lines were sustained for a longer duration and sequenced multiple times throughout the experiment. Consequently, rather than all mutants originating from a genetically identical ancestor, most mutants had a different direct ancestor. In addition, the fitness measure analyzed for replicate *j* of mutant line *i* was (*r*_*ij*_ *− r*_*wi*_)*/r*_*wi*_, rather than *r*_*ij*_, where *r*_*wi*_ is the maximum growth rate of the direct ancestor of line *i*. Growth rate was measured in triplicate for each line, but for the ancestral growth rate, *r*_*wi*_, we only had access to the average of the three measurements, not the individual measurements. We reanalyzed these data after generation scaling (i.e. *ln*(2)(*r*_*ij*_ *− r*_*wi*_)*/r*_*wi*_) using a mixed model with an intercept and random line effects only. In order to compare our between-line variance with that of Sane et al. (2018) we also fitted an equivalent model to the non-standardised growth rates (i.e. *r*_*ij*_). We also collated estimates of the mean selection coefficient in *E. coli* from three additional papers. Kibota and Lynch (1996) measured the fitness of wild-type MA lines and obtained an estimate of the lower bound for the mean of −0.012 (prior to a selection bias correction). Only a lower bound could be achieved because the lines were not sequenced and so a minimum mutation rate had to be estimated assuming the variance in selection coefficients was zero. Replacing the reported minimum number of mutations per genome per generation (1.7 *×* 10^−4^ - again, not corrected for selection bias) with the subsequent estimate of 1.014 *×* 10^−3^ from Lee et al. (2012), we obtain the estimate of the mean (rather than the lower bound) as −0.0014 after the *ln*(2) correction. Standard errors are not available. Baehr et al. (2025) obtained two estimates for a MutL strain - one for MA lines propagated in liquid culture and one for MA lines propagated on agar plates. After *ln*(2) scaling the estimates are(−5.8918*±*1.9408)*×*10^−4^ and (−5.4759*±*1.8715)*×*10^−4^, respectively.

Robert et al. (2018) generated single-cell MA lines using a MutH strain propagated in a ‘mother machine’ and after *ln*(2) scaling their estimate is −0.0021*±*0.0003.

## Results

A summary of parameter estimates, confidence intervals, and statistical significances are given in Table 2. A graphical depiction of the estimates is given in Figure 1. Using the expected fitness of the zero-mutant MA lines as the reference, we found that the genetically identical ancestor had a higher growth rate (*β*_*A*_=(2.736 *±* 0.534) *×* 10^−2^, P*<*0.001) and the average effect of a single mutation was to reduce the growth rate by a small and non-significant amount (*β*_*M*_ =(−2.584 *±* 13.730) *×* 10^−4^, P=0.851) (Table 2). After accounting for these mean effects, the between-line variance for single mutants 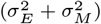 was estimated to be 3.208*×*10^−4^ and a value of zero could be clearly rejected 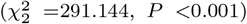,indicating systematic variation in fitness between lines. Notably, the between-line variance attributable to non-genetic effects 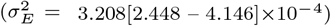 is considerably larger than the between-line variance attributable to genetic effects 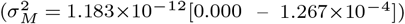 and removing the genetic effects 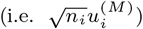 from the model did not produce a significant drop in model fit 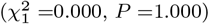 In contrast, removing the non-genetic effects 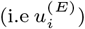 caused a significant drop in the log-likelihood 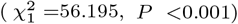. These findings suggest that systematic differences in fitness between-lines are mainly attributable to non-genetic effects and the between-line variance due to new mutations is small. Indeed, the mean and the variance of the DFE are both small in magnitude with *μ*_*s*_ =−1.397 [−19.795 – 17.313]*×*10^−4^ and 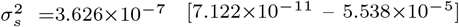 respectively. Restricting the analysis to non-synonymous mutations, INDELs in coding regions and large structural variants gave similar answers (*β*_*M*_ =(−1.719 *±* 2.022) *×* 10^−3^, P=0.399 and 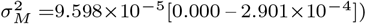. Because most studies lack zero-mutant MA lines, most studies use the ancestral line as a reference for calculating selection coefficients. Had we done this, our estimate of the mean and variance of the DFE would have been −20.308 [−37.084 – 3.302]*×*10^−3^ and 3.515*×*10^−7^ [6.87*×*10^−11^ – 5.2*×*10^−5^] respectively. Since the ancestor effect, *β*_*A*_, was comparable in magnitude to the between-line standard deviation of the non-genetic MA line effects (*β*_*A*_*/σ*_*E*_ = 1.528) it is possible that the ancestral effect is simply a non-genetic effect drawn from the same distribution of MA line non-genetic effects. Indeed, the ancestor effect was non-significant when a line effect (*u*^(*E*)^) was also fitted for the ancestor (*β*_*A*_=(2.736 *±* 1.869) *×* 10^−2^, P=0.146).

**Table 2.** Summary of growth rate analysis with estimates followed by their 95% confidence intervals and p-value.

| Parameter | Estimate | l-95% | u-95% | p-value |
| --- | --- | --- | --- | --- |
| $\beta_0$ | 1.088 | 1.070 | 1.106 | - |
| $\beta_A$ | 0.027 | 0.017 | 0.038 | $3.245 \times 10^{-7}$ |
| $\beta_M$ | $-2.584 \times 10^{-4}$ | $-2.944 \times 10^{-3}$ | $2.437 \times 10^{-3}$ | 0.851 |
| $\sigma_E^2$ | $3.208 \times 10^{-4}$ | $2.448 \times 10^{-4}$ | $4.146 \times 10^{-4}$ | $5.821 \times 10^{-14}$ |
| $\sigma_M^2$ | $1.183 \times 10^{-12}$ | 0.000 | $1.267 \times 10^{-4}$ | 1.000 |
| $\sigma_e^2$ | $1.172 \times 10^{-3}$ | $1.103 \times 10^{-3}$ | $1.245 \times 10^{-3}$ | - |

**Figure 1.**
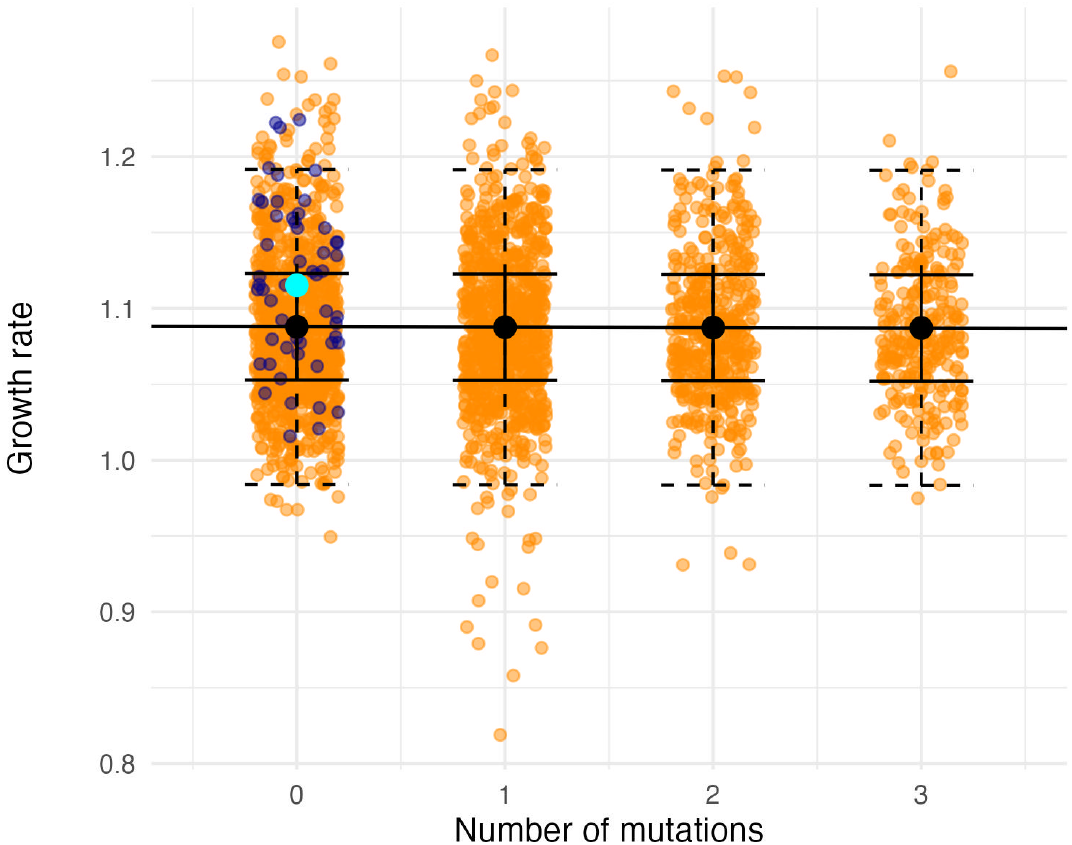
Maximum growth rate (log(OD) per hour) with replicates of MA lines in orange and replicates of the ancestral line in blue. The solid black line is the fitted regression with black dots representing the predicted mean fitness for lines with different numbers of mutation. The whiskers indicate the lower and upper bound of the 95% quantiles for the line effects (full black bars) and the observations (dashed black bars). The cyan point is the estimated mean growth rate of the ancestral line.

Reanalysis of data from Sane et al. (2018) gave estimates for the mean and variance of the DFE of 0.031*±*0.012 and 0.011 [7.968*×*10^−3^-0.016], respectively. The between-line variance in the raw growth rate 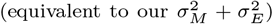 was estimated to be 0.023 [0.016-0.032], considerably larger than our estimate of the total between-line variance (3.208*×*10^−4^). Estimates of the mean selection coefficient from our study and the literature are summarised in Figure 2.

**Figure 2.**
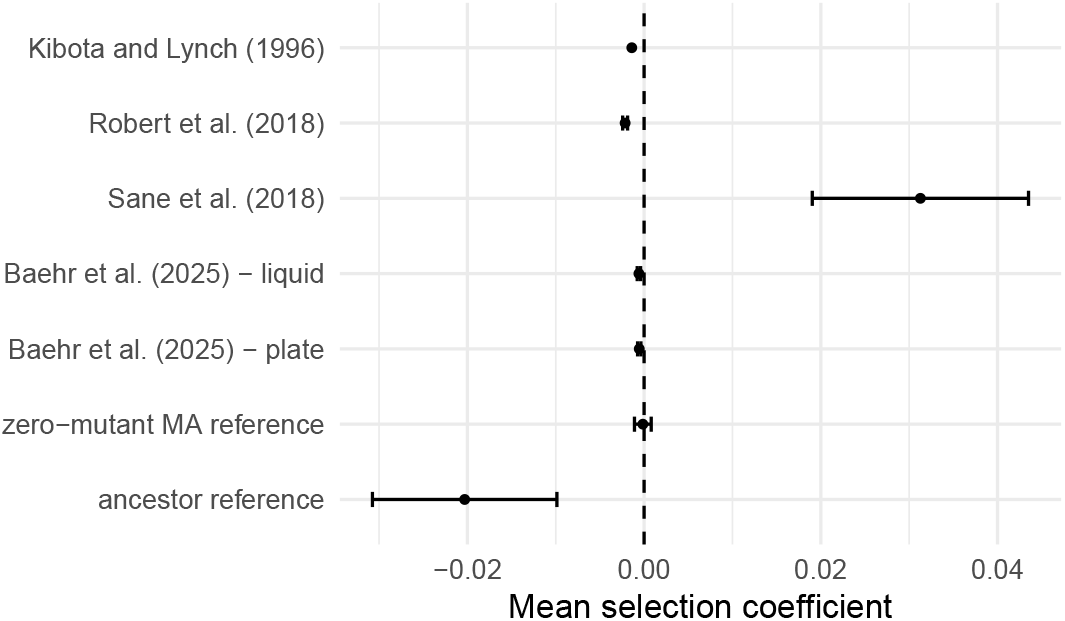
Estimates *±* standard error of the mean of the distribution of fitness effects (DFE) of new mutations in *E. coli*. Kibota and Lynch (1996) gave a lower bound for the mean of the DFE without standard errors: the point estimate was turned into an estimate of the mean (rather than a lower bound) using the subsequently published mutation rate of 1.014 *×* 10^−3^ mutations per genome per generation (Lee et al., 2012). Robert et al. (2018) used a MutH strain and Baehr et al. (2025) a MutL strain. All published estimates were scaled by *ln*(2) to place them in units of generations (Chevin, 2011).

## Discussion

In this study, we attempted to estimate the distribution of fitness effects (DFE) using fitness measurements of *E. coli* lines that had accumulated single mutations under mutation accumulation (MA).

Previous attempts at directly measuring the DFE have used multi-mutant lines, yet these can only give precise information about the mean and variance of the DFE, whereas higher moments are hard to estimate (Keightley, 1994). However, despite high replication, we could not reject a DFE with a point mass at zero, and can only put a lower bound (at 95% credibility) on the mean fitness effect of −1.98*×*10^−3^. The power to estimate the DFE was compromised by the fitness effects of mutations being much smaller in magnitude than non-genetic causes of fitness differences between lines. Previous work in microbes has assumed these non-genetic effects to be negligibly small. Below we provide evidence that non-genetic fitness effects are a prevalent problem in microbial MA lines, causing fitness effects to be incorrectly estimated. We go on to give suggestions for alternative protocols that can be used to get more precise DFE estimates that are not confounded by non-genetic effects.

Previous estimates of the DFE in microbes have neither quantified nor controlled for sources of between-line variance that are not due to new mutations. Consequently, it is hard to determine whether the large non-genetic between-line variance we observe is specific to our experimental set-up or a more widespread problem. To investigate the prevalence of these effects, we compared the magnitude of our between-line variance in fitness to one previously reported from single-mutant MA lines accumulated in *E. coli* (Sane et al., 2018). While it was not possible to differentiate between genetic and non-genetic sources of between-line fitness under this experimental design, we would expect the between-line variance in fitness to have a similar magnitude as our *genetic* between-line fitness variance if non-genetic sources of variation were absent. In contrast, the between-line variance from Sane et al. (2018) far exceeded not only our genetic between-line variance, but also our total between-line variance. This result suggests that the degree of bias under certain experimental protocols may even exceed the degree of bias we observe. In this instance, the greater non-genetic between-line variance might be due to the fitness assay procedure, where Sane et al.’s (2018) replicates were taken from the same culture, whereas our replicates were re-cultured each time from a frozen stock. This may have been compounded by replicate fitness assays appearing sequentially on the same plate, whereas our randomisation procedure should have eliminated any plate/positional effects from the between-line variance. Alternatively, the greater between-line variance in Sane et al. (2018) may be statistical in origin: given that the same ancestral growth rate measurement was used when calculating a mutation’s relative fitness for all three replicates, ancestral measurement error may be contributing to the large between-line differences. Our estimate of the mean selection coefficient was small (−1.397*×*10^−4^), but largely consistent with estimates from multi-mutant MA lines (see Figure 2). However, had we used the ancestral line as a reference, rather than the zero-mutant MA lines derived from it, our estimate of the mean selection coefficient (−0.02) would have exceeded in magnitude these previous estimates by some margin. Our reanalysis of Sane et al. (2018) estimated the average selection coefficient to be 0.031. Given this is opposite in sign to expectation, and of a magnitude more consistent with selection coefficients of gene knock-outs using transposon insertions (~ −0.03 Couce et al. (2024); Elena et al. (1998)), it seems likely that the ancestral lines were also subject to some environmental perturbation as in our study.

Although the use of single-mutant lines is optimal for preserving information about the complete DFE, rather than just its mean and variance, the method comes with the caveat that genetic between-line effects will be small in magnitude compared to the non-genetic between-line and within-line fitness effects. This will mean that estimates of the DFE from single-mutant lines will be more biased by non-genetic effects if they are unaccounted for, and will be subject to higher sampling error even when they are accounted for. Indeed, even with 14 replicates for nearly 200 lines we failed to detect significant genetic between-line effects. While increasing the number of lines and replicates would afford more power, a different experimental setup may be preferable. For example, competitive assays, where single mutants are competed directly against a reference strain would effectively reduce the within-line variance since differences in fitness between the mutant and reference strain can be calculated within wells, rather than between. Gallet et al. (2012) showed that selection coefficients as small as 10^−3^ can be estimated by utilizing flow cytometry to distinguish fluorescently labelled mutant and reference lines (Heilbron et al., 2014). However, estimating such small fitness differences accurately requires that misclassification rates are essentially zero. Therefore, we recommend performing MA on a fluorescently-labelled ancestral strain which can easily be distinguished from a complementary labelled reference strain. Similar competitive fitness assays have also been done using bulk sequencing, where barcoded mutants are pooled and their relative frequencies measured (Ardell et al., 2024; Couce et al., 2024). While bulk sequencing could also be used to estimate frequency changes of spontaneous mutations, the sequencing depth required to get precise estimates might be prohibitive. A more cost-effective approach would be to conduct MA in parallel on barcoded ancestral lines, with barcode effects controlled for by estimating their fitness effects from competitive assays of the ancestral lines prior to MA. Alternatively, a promising possibility would be to use a mother-machine, where cell lineages are individually propagated in capillary tubes and differences in growth rate between mother and daughter cells can be measured (Wang et al., 2010). Coupling this information directly with data on the occurrence, or not, of mutations would allow estimates of the DFE (Robert et al., 2018). However, harvesting lineages for sequencing and/or running experiments on the timescale of weeks or months is not yet feasible, putting the study of spontaneous mutations currently out of reach.

The protocols recommended above envisage having access to lines without accumulated mutations to quantify non-genetic between-line variance. However, in many experiments, zero mutant lines are not available due to high genomic mutation rates or MA lines running for many generations. While the relative bias caused by non-genetic effects is reduced with multi-mutant lines, it may still be necessary to minimize, determine and correct for them. One possibility would be to propagate replicates from the same MA line for extended periods prior to fitness measurements in the hope that any shared environmental effect from their common ancestor has dissipated (Lynch, 1985). For example, in Vassilieva and Lynch (1999), *C. elegans* fitness measurements were taken from third-generation progeny to ensure that maternal and grandmaternal effects did not contribute to the between-line variance. For *E. coli* grown in a mother-machine, environmental effects were estimated to decay over approximately ten generations (Supporting Information in Robert et al., 2018), consistent with the phenotypic memory conferred by protein transmission (Lambert and Kussel, 2014) or correlation of sister-cell cell-cycle time and cell size (Vashistha et al., 2021). This time scale seems to contradict our result that differences between lines due to non-genetic effects still persist in our study despite replicates of the same line being separated by approximately 50 generations. However, we stress that although the evidence for shared environmental effects in our experiment is strong, the effects themselves are small in magnitude and would likely go undetected without substantial replication. In addition, in our MA set-up, lines have been propagated independently for approximately 1,000 generations allowing them to accumulate environmental differences that may be greater than those observed in studies such as Robert et al. (2018) and Vashistha et al. (2021). Moreover, several mechanisms are known to underpin the transfer of environmental effects over extended periods such as methylation status (Hernday et al., 2002), self-templating proteins (Yuan and Hochschild, 2017) and self-sustaining transcriptional feedbacks (Gardner et al., 2000) although examples are limited, and often synthetic.

An alternative to extended propagation of replicates prior to fitness measurement is to regress the mean and between-line variance in fitness on the number of mutations accumulated. Robust estimates of the mean and variance of the DFE are then possible if non-genetic sources of between-line variance are independent of mutation number. This is likely to be the case if variation in mutation number is simply due to stochasticity in mutation number from MA lines ran for the same number of generations. However, to generate variation in mutation number it is common to sample MA lines at different generations (e.g. Chebib et al., 2024) which can induce a relationship between the number of mutations and the effects of non-mutational processes. Of relevance here is when MA lines are so short that the variance caused by inherited non-genetic effects has yet to equilibrate at its stationary variance, and is therefore still increasing in time. While this seems unlikely in our set up where MA lines have been run for approximately 1,000 generations, this may prove problematic for shorter MA experiments on larger organisms or microbes with artificially high mutation rates. In addition, for sexual organisms, if the residual heterozygosity in the MA ancestor exceeds the expected equilibrium heterozygosity in the MA lines, there will be a period of accelerated divergence in early generations (Lynch and Hill, 1986). Similarly, natural selection against new mutations will slow the rate of divergence in later generations as mutations get purged (Mackay et al., 1995). While all of these processes may generate a spurious relationship between mutation number and the between-line variance, and as a result generate biased estimates of the DFE (see SI for more detail, including theory), regressing the between-line variance in fitness on the number of mutations is both simple and likely results in better estimates than simply ignoring non-genetic sources of between-line variance.

To summarise, in this study we have demonstrated that the selection coefficients of spontaneous single mutations in *E. coli* are on average only slightly deleterious, consistent with estimates from most other studies (Figure 2). While this conclusion seems to contradict the widely held assumption that mutations are on average considerably more deleterious, it has usually not been recognised that the estimate from Kibota and Lynch (1996) is a lower bound and largely consistent with more recent estimates (Robert et al., 2018; Baehr et al., 2025). Although correcting these estimates for selection bias is likely to produce revised estimates that are more negative (Wahl and Agashe, 2022), very large shifts are not expected when fitness effects are small (Grosse-Sommer and Hadfield, 2026). Our unique experimental design also allowed us to demonstrate non-genetic fitness effects between lines, which severely bias estimates of the DFE if they are not accounted for.

## Supporting information

Fitness Analyses

Mutation Calling

Antedependence Model

## Data availability

The raw sequence data from our E. coli MA experiment has been uploaded to the NCBI Sequence Read Archive under BioProject ID PRJNA1466858. All data and code used in the analyses can be found at https://github.com/JGrosseSommer/Ecoli_MA.

## Acknowledgements

We thank Mrudula Sane and Deepa Agashe for sharing data and extensive discussion. We thank Rob Ness and Jobran Chebib for help and insight. This work was supported by funding from NERC through an E4 DTP studentship.

