## Supplementary material for "Fitness effects of new mutations are small and heavily confounded with non-genetic sources of variation in Escherichia coli": Fitness Analyses: Fitness_Analyses.html

Esherichia coli Fitness Analysis


### *Esherichia coli* Fitness Analysis

- 1 Data
- 2 Data filtering
  - 2.1 High starting OD
  - 2.2 Empty wells
  - 2.3 Extra wells
  - 2.4 Filtering
- 3 Growth Rate and Meta Data
  - 3.1 Empty wells
- 4 Models
  - 4.1 Main Model
  - 4.2 Obtaining the DFE
    - 4.2.1 Estimating the DFE using the ancestor as a reference
  - 4.3 Is the ancestor effect comparable to other zero-mutant line effects?
  - 4.4 Mutation type
  - 4.5 Significance testing of random effect variances
    - 4.5.1 Genetic line effects
    - 4.5.2 Environmental line effects
- 5 Comparison with Sane, Miranda, and Agashe (2018)

220 *Escherichia coli* lines underwent 36 days of mutation accumulation (MA), the time it takes to accumulate one mutation on average. The aim of the study was to estimate the distribution of fitness effects of new mutations (DFE) by measuring the growth rate of 188 of these lines and their common ancestor. Since the protocol also generated many lines that had accumulated no mutants, we could also estimate the distribution of fitness effects across genetically identical MA lines. We refer to this as the distribution of fitness effects due to environmental effects (EDFE) although acknowledge that we use the word ‘environmental’ to capture all systematic differences between lines not due to DNA mutations.

### 1 Data

The 188 *Escherichia coli* mutation accumulation lines and four ancestral replicate lines were split into two sets, each containing 94 MA lines and 2 replicates of the ancestral line. For each set, the frozen lines (`line`) were grown while shaking for 16 hours at 37\(^{\circ}\)C in 10ml LB media, after which 10\(\mu\)L of culture was transferred to a 96-well filled with 190\(\mu\)L of LB. Optical density measurements (`OD`) were taken every 5 minutes for four hours (`timepoint` in hours). This was repeated 14 times, and the membership of each set and the position of each line (`row` and `column`) on the `plate` was randomised between replicates. The `date` on which the lines were defrosted and grown is recorded together with the `person` (JDH or DN) inoculating the cultures. JGS was responsible for inoculating all 96 well plates.

```
library(dplyr)
library(gcplyr)
library(ggplot2)
library(tidyr)

od_df<-read.csv(file.path(root, "meta/growth_data.csv"))
```

We can plot the optical density (OD) over time (with time jittered):

```
plot(OD~jitter(timepoint,2), data=od_df, pch=16, cex=0.2, bty="l", ylab="Optical Density", xlab="Time (hours)")
```

While the bulk of the growth curves look reasonable, there are clearly some issues with no growth in some wells and some unusually high starting OD’s.

### 2 Data filtering

#### 2.1 High starting OD

There are 38 replicates that have a starting OD higher than 0.2.

```
od_df <- mutate(group_by(od_df, plate, row, column), highOD = OD[timepoint==0.08]>0.2)
tapply(od_df$highOD, od_df$plate, sum)/48
```

```
##  1  2  3  4  5  6  7  8  9 10 11 12 13 14 15 16 17 18 19 20 21 22 23 24 25 26 
##  1  1  0  0  1  0  1  0  0  0  0  0  0  0  0  0  0  0  0  0  0 16 15  0  0  0 
## 27 28 
##  0  3
```

31 of these replicates were on plates 22 and 23. The function `plot_plate` allows us to visualise data in plate layout:

```
plot_plate <- function(df, z, palette = "plasma", digits = 2) {

  ggplot(df, aes(x = as.factor(column), y = factor(row, levels=rev(LETTERS[1:8])), fill = df[[z]])) +
    geom_tile(colour = "grey50") +
    geom_text(aes(label = round(df[[z]], digits)),
              size = 3, colour = "white") +
    scale_fill_viridis_c(option = palette, na.value = "white") +
    coord_equal() +
    labs(x = "Column", y = "Row", fill = z) +
    theme_minimal(base_size = 14) +
    theme(panel.grid = element_blank())
}
```

We can see that wells with unusually high starting OD are clustered on the two plates:

```
plot_plate(subset(od_df, timepoint==0.08 & plate==22), z = "OD")
```

```
plot_plate(subset(od_df, timepoint==0.08 & plate==23), z = "OD")
```

This does not look like systematic pipetting error, but rather some local issue with the plates. Of the remaining seven wells with high starting OD, two were known to be due to pipetting error and were doubly inoculated (5-H-1 and 28-G-1 in red) and two were unintentionally blacked-out (2-A-8 and 1-A-5 in blue, see below)

```
plot(OD~jitter(timepoint,2), data=od_df, pch=16, cex=0.2, bty="l", ylab="Optical Density", xlab="Time (hours)")
points(OD~timepoint, data=subset(od_df, plate==5 & row=="H" & column==1), pch=16, cex=0.5, col="red")
points(OD~timepoint, data=subset(od_df, plate==28 & row=="G" & column==1), pch=16, cex=0.5, col="red")
points(OD~timepoint, data=subset(od_df, plate==2 & row=="A" & column==8), pch=16, cex=0.5, col="blue")
points(OD~timepoint, data=subset(od_df, plate==1 & row=="A" & column==5), pch=16, cex=0.5, col="blue")
```

For the remaining three wells (7-B-6, 28-B-4, 28-F-7) the starting OD was only a little above 0.2 and the growth curves look fine:

```
plot(OD~timepoint, data=subset(od_df, highOD & plate%in%c(7,28) & row!="G"), pch=16, cex=0.5, bty="l", ylab="Optical Density", xlab="Time (hours)")
```

In order to retain these growth curves for later we will change their highOD flag to FALSE:

```
with(od_df, highOD[which(highOD & plate%in%c(7,28) & row!="G")]<-FALSE)
```

#### 2.2 Empty wells

In 40 cases the initial culture looked like it hadn’t grown, and in an error of judgement the culture was not assayed for growth and the well was either blacked-out (2-A-8 and 1-A-5, see above) or left empty (otherwise).

```
plate.blank<-c(1, 2, 3, 5, 7, 9, 11, 11, 11, 15, 15, 15, 17, 19, 19, 21, 21, 21, 23, 23, 23, 25, 25, 27, 2, 8, 12, 18, 20, 22, 22, 22, 24, 24, 26, 28, 28, 28, 10, 10)
row.blank<-c("A", "A", "A", "B", "D", "H", "B", "D", "F", "F", "G", "H", "G", "B", "C", "A", "E", "F", "E", "F", "F", "C", "C", "D", "A", "C", "B", "F", "H", "A", "B", "H", "G", "F", "F", "C", "A", "E", "D", "H")
column.blank<-c(5, 8, 6, 3, 10, 5, 2, 3, 7, 6, 6, 12, 7, 12, 12, 6, 7, 8, 2, 2, 12, 2, 8, 2, 8, 12, 8, 4, 5, 8, 10, 10, 1, 1, 1, 2, 9, 9, 8, 12)
```

```
od_df <- mutate(group_by(od_df, plate, row, column), empty = paste(plate, row, column)%in%paste(plate.blank, row.blank, column.blank))
```

Note four (rather than two) wells do not appear empty - the additional wells are 22-H-10 and 23-F-2 for which there appeared to be an issue with the plate (see above). The number of empty wells per line is clearly overdispersed:

```
n_empty<-tapply(od_df$empty, od_df$line, sum)/48
hist(n_empty, breaks=100, ylab="Number of Lines", xlab="Number of Empty Wells")
```

In particular line 180 repeatedly failed to grow (10 times out of 14).

```
n_empty[which(n_empty!=0)]
```

```
## 118  16 180 203  29  43  68  81  84  97 
##   5   6  10   1   1   4   2   7   2   1
```

#### 2.3 Extra wells

In addition, well 5-F-9 also appears empty. This replicate was of line 180 which repeatedly failed to grow (see above).

```
od_df <- mutate(group_by(od_df, plate, row, column), extra_empty = OD[timepoint==2]<0.2 & !empty)
plot(OD~jitter(timepoint,2), data=od_df, pch=16, cex=0.2, bty="l", ylab="Optical Density", xlab="Time (hours)")
points(OD~timepoint, data=subset(od_df, extra_empty), pch=16, cex=0.5, col="red")
```

Nevertheless, three replicates of line 180 did grow despite starting at a low OD:

#### 2.4 Filtering

In what follows we remove empty wells and those in which the starting OD was greater than 0.2 (except 7-B-6, 28-B-4 and 28-F-7 which look fine):

```
od_df <- subset(od_df, !highOD & !empty & !extra_empty)
```

```
plot(OD~jitter(timepoint,2),data=od_df, pch=16, cex=0.2, bty="l", ylab="Optical Density", xlab="Time (hours)")
```

### 3 Growth Rate and Meta Data

In order to get an estimate of the growth rate in the exponential phase we can use gcplyr (Blazanin 2024). The change in log OD is computed over consecutive time-points for each well and a rolling 3-timepoint average calculated:

```
df <- mutate(group_by(od_df, plate, row, column), deriv = calc_deriv(x = timepoint, y = OD, percapita = TRUE, blank = 0, window_width_n = 3, trans_y = 'log'))
```

For each well, the maximum average change is taken as the growth rate:

```
df <- summarize(df, gr = max_gc(deriv), across(-c(timepoint, OD, deriv),first))
```

Since growth rates may be different in the inner wells, we can create a factor for modelling this

```
df <- mutate(df, inner = as.numeric(!(row %in% c('A', 'H') | column %in% c(1, 12))))
```

We can also add the number of mutations per line

```
mutations<-read.csv(file.path(root, "meta/real_mutations.csv"))
```

```
nmutations<-table(factor(mutations$line, levels=unique(df$line)))
df$mutation<-as.numeric(nmutations[df$line])
```

and have two indicator variables for whether a line is the ancestor (`anc`) or not (`noanc`):

```
df$anc<-as.numeric(df$line=="anc")
df$noanc<-as.numeric(df$line!="anc")
```

A violin plot of the growth rates reveals an outlier with a very high growth rate:

```
ggplot(data=df,aes(x='',y=gr)) + geom_violin() + geom_boxplot(width=0.1)+scale_y_continuous(name="growth rate")
```

It seems that the high growth rate is driven by the anomalous OD readings (in red)

```
plot(log(OD)~timepoint, data=subset(od_df, plate==23 & row=="H" & column==1), ylab="log(OD)", xlab="Time (hours)")
points(log(OD)~timepoint, data=subset(od_df, plate==23 & row=="H" & column==1 & timepoint>1.55 & timepoint<1.7), col="red", pch=16, cex=1.25)
```

Consequently, we also remove this data point:

```
df<-subset(df, gr<1.5)
```

Line 95 accumulated a mutation in the mismatch-repair gene MutL and consequently had an elevated mutation rate with 17 fixed mutations observed. There is evidence from the sequencing data that an addition five mutations arose after mutation accumulation but prior to sequencing. Given new mutations are also likely to arise during the growth assay, and data from this line are likely to have high leverage (all other lines had between zero and five), we removed this line from the analysis:

```
df<-subset(df, line!=95)
```

#### 3.1 Empty wells

40 replicates were not assayed because the initial culture looked like it hadn’t grown, with certain lines being overrepresented. If the initial culture did not grow because of the accumulated mutations in the line, this will bias our estimates of the DFE. However, if we compare the growth rates of lines where at least one replicates was excluded with the growth rates of lines where no replicates were excluded the difference is small and non-significant:

```
nz_empty<-n_empty[which(n_empty!=0)]
summary(lm(gr~as.factor(line%in%nz_empty), data=df))
```

```
## 
## Call:
## lm(formula = gr ~ as.factor(line %in% nz_empty), data = df)
## 
## Residuals:
##       Min        1Q    Median        3Q       Max 
## -0.268870 -0.035068 -0.004517  0.033388  0.187799 
## 
## Coefficients:
##                                    Estimate Std. Error  t value Pr(>|t|)    
## (Intercept)                        1.087797   0.001040 1045.760   <2e-16 ***
## as.factor(line %in% nz_empty)TRUE -0.008608   0.006338   -1.358    0.175    
## ---
## Signif. codes:  0 '***' 0.001 '**' 0.01 '*' 0.05 '.' 0.1 ' ' 1
## 
## Residual standard error: 0.05231 on 2597 degrees of freedom
## Multiple R-squared:  0.0007098,  Adjusted R-squared:  0.000325 
## F-statistic: 1.845 on 1 and 2597 DF,  p-value: 0.1745
```

Indeed, if we plot the growth rates of the four lines with the most replicates excluded (the coloured points) they do not obviously grow more slowly than other lines:

```
boxplot(gr~as.factor(line%in%nz_empty), data=df, ylab="Growth Rate", xlab="At least one replicate excluded")

worst_four<-subset(df, line%in%names(nz_empty)[which(nz_empty>4)])

points(gr~I(2+(as.numeric(as.factor(line))-2.5)/6), col=as.numeric(as.factor(line)), data=worst_four)
```

and the line (in red) with the lowest average growth rate actually has no mutations:

```
with(worst_four, mutation[match(levels(as.factor(line)), line)])
```

```
## [1] 1 0 1 1
```

### 4 Models

#### 4.1 Main Model

We fit a linear mixed model to the growth rates with the aim of estimating the mean and variance of the distribution of fitness effects of new mutations (DFE). We can divide the terms of the model into those that are biologically interesting:

- `(Intercept)` (fixed): expected growth rate of a zero-mutant line.
- `mutation` (fixed): the linear change in fitness per mutation, related to the mean of the DFE.
- `anc` (fixed) effect of a line being an ancestor.
- `(sqrt(mutation)-1|line)` (random) the effect of each mutation in a specific non-ancestral line. The variance of these effects is related to the variance of the DFE.
- `(noanc-1|line)` (random) the effect of being a specific non-ancestral line when the number of new mutations is zero. The variance of these effects is the variance of the EDFE.

and those that control for technical variation:

- `person` (fixed) effect of the person inoculating the cultures.
- `inner` (fixed) effect of whether growth rates are different in inner wells.
- `(1|plate)` (random) effect of being on a specific plate.
- `(1|plate:column)` (random) effect of being in a specific column of a specific plate.
- `(1|plate:row)` (random) effect of being in a specific row of a specific plate.

```
library(lmerTest)
library(MCMCglmm)

m1<-lmer(gr~mutation + anc + person + inner + (noanc-1|line) + (sqrt(mutation)-1|line) + (1|plate) + (1|plate:column) + (1|plate:row), data=df)
summary(m1)
```

```
## Linear mixed model fit by REML. t-tests use Satterthwaite's method [
## lmerModLmerTest]
## Formula: 
## gr ~ mutation + anc + person + inner + (noanc - 1 | line) + (sqrt(mutation) -  
##     1 | line) + (1 | plate) + (1 | plate:column) + (1 | plate:row)
##    Data: df
## 
## REML criterion at convergence: -9436
## 
## Scaled residuals: 
##     Min      1Q  Median      3Q     Max 
## -4.7550 -0.6015 -0.0610  0.5026  3.7585 
## 
## Random effects:
##  Groups       Name           Variance  Std.Dev. 
##  plate.column (Intercept)    4.954e-05 7.038e-03
##  plate.row    (Intercept)    1.815e-04 1.347e-02
##  line         sqrt(mutation) 1.183e-12 1.088e-06
##  line.1       noanc          3.208e-04 1.791e-02
##  plate        (Intercept)    1.083e-03 3.291e-02
##  Residual                    1.172e-03 3.423e-02
## Number of obs: 2599, groups:  
## plate:column, 336; plate:row, 224; line, 188; plate, 28
## 
## Fixed effects:
##               Estimate Std. Error         df t value Pr(>|t|)    
## (Intercept)  1.088e+00  9.252e-03  2.961e+01 117.575  < 2e-16 ***
## mutation    -2.584e-04  1.373e-03  1.846e+02  -0.188  0.85090    
## anc          2.736e-02  5.339e-03  2.261e+03   5.124 3.24e-07 ***
## personJDH    6.857e-03  1.267e-02  2.603e+01   0.541  0.59293    
## inner       -5.776e-03  1.899e-03  7.404e+02  -3.041  0.00244 ** 
## ---
## Signif. codes:  0 '***' 0.001 '**' 0.01 '*' 0.05 '.' 0.1 ' ' 1
## 
## Correlation of Fixed Effects:
##           (Intr) mutatn anc    prsJDH
## mutation  -0.162                     
## anc       -0.093  0.276              
## personJDH -0.685  0.003  0.000       
## inner     -0.129 -0.001  0.027  0.000
## optimizer (nloptwrap) convergence code: 0 (OK)
## boundary (singular) fit: see help('isSingular')
```

The biological findings of this model are:

- Assuming no cell death, we can divide \(ln(2)\) by the intercept to get an estimate of the generation time of a mutant-free line (grown in an outer well and inoculated by DN): 38 minutes.
- The `mutation` effect is the change in log growth rate caused by a single mutation - it is small and not significantly different from zero.
- The `anc` effect is positive suggesting that the ancestral line grows faster than the average growth rate of the mutant-free lines.
- The `line.1` (`noanc`) variance, which is due to between-line differences not caused by mutations, explains approximately 11% of the total variance (after controlling for the fixed effects).
- The estimate of the `line` (`sqrt(mutation)`) variance is bound at zero and suggests that single mutations generate very little, if any, between-line variance.

The diagnostic plot of the residuals against the fitted values looks OK:

```
plot(m1)
```

We can also plot the data and some model predictions in order to get a feel for what the model is telling us

```
intercept=fixef(m1)['(Intercept)']
mutation_effect=fixef(m1)['mutation']
ancestor_effect=fixef(m1)['anc']

env_variance<-as.numeric(VarCorr(m1)[['line.1']])
tech_variance<-as.numeric(VarCorr(m1)[['plate.column']]+VarCorr(m1)[['plate.row']]+VarCorr(m1)[['plate']])
mut_variance<-as.numeric(VarCorr(m1)[['line']])
res_variance<-sigma(m1)^2

pred_mutations<-0:3

pred_mean<-intercept+mutation_effect*pred_mutations
pred_var<-env_variance+res_variance+mut_variance*pred_mutations+tech_variance
pred_bl_var<-env_variance+mut_variance*pred_mutations

pred_ancestor<-intercept+ancestor_effect

pred_lower_q<-qnorm(0.025,pred_mean,sqrt(pred_var))
pred_upper_q<-qnorm(0.975,pred_mean,sqrt(pred_var))

pred_lower_q_bl<-qnorm(0.025,pred_mean,sqrt(pred_bl_var))
pred_upper_q_bl<-qnorm(0.975,pred_mean,sqrt(pred_bl_var))

plot_data<-cbind(pred_mutations,pred_mean,pred_lower_q,pred_upper_q,pred_lower_q_bl,pred_upper_q_bl)

df_plot <- df %>% 
  mutate(point_col = case_when(
    noanc == 0                          ~ "blue",
    mutation <= 3 & noanc == 1          ~ "orange",
    TRUE                                ~ "other")
  )

df_plot<-df_plot[order(df_plot$point_col, decreasing=TRUE),]

CI <- ggplot(plot_data, aes(pred_mutations, pred_mean)) +

  ## a) raw-data points (jittered) -------------
  geom_jitter(data = filter(df_plot, point_col != "other"),
              aes(x = mutation, y = gr, colour = point_col),
              width  = .20, height = 0, alpha = .5, show.legend = FALSE) +

  scale_colour_manual(values = c(orange = "darkorange",
                                 blue   = "darkblue")) +

  ## b) regression line ------------------------
  geom_abline(intercept = intercept, slope = mutation_effect) +

  ## c) fitted means & error bars --------------
  geom_point(size = 3) +
  geom_errorbar(aes(ymin = pred_upper_q, ymax = pred_lower_q),
                width = .5, colour = "black", linetype = "dashed") +
  geom_errorbar(aes(ymin = pred_upper_q_bl, ymax = pred_lower_q_bl),
                width = .5, colour = "black") +

  ## d) ancestor point -------------------------
  annotate("point", x = 0, y = pred_ancestor, colour = "cyan", size = 3, alpha = 1)+

  ## e) axes, labels, limits -------------------
  scale_x_continuous(limits = c(-.5, 3.5),
                     name   = "Number of mutations") +
  ylab("Growth rate") +

  ## f) theme ----------------------------------
  theme_minimal(base_size = 11) +
  theme(
 #   axis.text   = element_text(face = "bold"),
   axis.title  = element_text(size = 11),
    axis.title.y = element_text(margin = margin(t = 10, r = 20, b = 20))
  )

  plot(CI)
```

(#fig:main\_plot)Raw growth rate (log(OD) per hour) for lines grouped by number of accumulated mutations. Replicates of non-ancestral lines are in orange and replicates of ancestral lines are in blue. The solid black line is the fitted regression with black dots representing the predicted mean fitness for lines with different numbers of mutation. The whiskers indicate the 95% quantiles for the line effects (full black bars) and the observations (dashed black bars). The cyan data point is the estimated mean ancestral growth rate.

```
  if(save){
  ggsave(file.path(root,'models/fitness.jpeg'),width = 5, height = 4,units='in')
  }
```

There seems to be quite a few reduced growth rates in the single mutant class - many of these are associated with line 45 which has a frameshift mutation in the *SspA* gene:

```
table(subset(df, gr<0.95)$line)
```

```
## 
## 103 115 117  18 218  45  98 
##   1   2   1   2   1  10   1
```

We can highlight the data points in this line in red.

```
df_45 <- df_plot %>%filter(line == 45)
CI2 <- CI +                                          # start from CI
  geom_jitter(data = df_45,                   # only these rows
              aes(x = mutation, y = gr),             # same aesthetics
              width  = .20, height = 0,
              colour = "red",                        # fixed colour
              size   = 2.2, alpha = .8,
              show.legend = FALSE)
  plot(CI2)
```

Figure 4.1: Raw growth rate (log(OD) per hour) for lines grouped by number of accumulated mutations. Replicates of non-ancestral lines are in orange and replicates of ancestral lines are in blue. The solid black line is the fitted regression with black dots representing the predicted mean fitness for lines with different numbers of mutation. The whiskers indicate the 95% quantiles for the line effects (full black bars) and the observations (dashed black bars). The cyan data point is the estimated mean ancestral growth rate and the red points are from line 45

```
   if(save){
  ggsave(file.path(root,'models/fitness2.jpeg'),width = 5, height = 4,units='in')
  }
```

#### 4.2 Obtaining the DFE

The previous model provides a model for the growth rates of each line. However,
to obtain the DFE we need to compute the distribution of relative fitnesses for single-mutant lines compared to zero-mutant lines. The expected growth rate of single-mutant lines (\(r\_w\)) is the intercept which we denote \(\beta\_0\).

\[\begin{equation}
r\_w = \beta\_0
\end{equation}\]

The expected growth rate of line \(i\) is

\[\begin{equation}
r\_i = \beta\_0 + n\_i\beta\_M+\sqrt{n\_i}u^{(M)}\_i
\end{equation}\]

where the expectation is taken over the distribution of environmental effects. \(n\_i\) is the number of mutations line \(i\) carries (`mutation`) and \(\beta\_M\) is the corresponding slope. \(u^{(M)}\_i\) is the line slope as parameterised in the term `(sqrt(mutation)|line)`.

The selection coefficient for MA line \(i\) is \(r\_i-r\_w\), although this is in units *per hour* and is therefore hard to compare across studies and species (See Equation 2.2 in Chevin (2011) and Equation 7 in Fink and Manhart (2024)). To make selection coefficients more comparable, they can be put on the scale of generations by using Equation 3.2 in Chevin (2011):

\[\begin{equation}
s\_i=\frac{r\_i-r\_w}{r\_w}ln(2)
\end{equation}\]

Setting \(n=1\), the distribution of generation-scaled fitness effects (the DFE) from our model is normal with mean

\[\begin{equation}
\mu\_s = \frac{\beta\_M}{\beta\_0}ln(2)
\end{equation}\]

and variance

\[\begin{equation}
\sigma^2\_s = \frac{\sigma^2\_M}{\beta\_0^2}ln(2)^2
\end{equation}\]

Since the mean and the variance of the DFE are functions of model parameters we refit model `m1` using Markov chain Monte Carlo in order to obtain credible intervals.

```
m1_mcmc<-MCMCglmm(gr~mutation+anc+person+inner, 
                  random=~line+idh(sqrt(mutation)):line+plate+plate:column+plate:row, 
                  data=as.data.frame(df), 
                  prior=list(R=IW(1, 0.002), G=F(1,1000)), 
                  longer=10, verbose=FALSE)
```

The model estimates are very close to those obtained using restricted maximum likelihood:

```
summary(m1_mcmc)
```

```
## 
##  Iterations = 30001:129901
##  Thinning interval  = 100
##  Sample size  = 1000 
## 
##  DIC: -9789.267 
## 
##  G-structure:  ~line
## 
##      post.mean l-95% CI  u-95% CI eff.samp
## line 0.0002977 0.000178 0.0004013    923.4
## 
##                ~idh(sqrt(mutation)):line
## 
##                     post.mean  l-95% CI  u-95% CI eff.samp
## sqrt(mutation).line 3.661e-05 1.729e-10 0.0001363    857.7
## 
##                ~plate
## 
##       post.mean  l-95% CI u-95% CI eff.samp
## plate  0.001219 0.0006559 0.001989    892.1
## 
##                ~plate:column
## 
##              post.mean  l-95% CI  u-95% CI eff.samp
## plate:column 4.881e-05 1.121e-05 8.086e-05     1000
## 
##                ~plate:row
## 
##           post.mean  l-95% CI  u-95% CI eff.samp
## plate:row 0.0001853 0.0001314 0.0002454     1000
## 
##  R-structure:  ~units
## 
##       post.mean l-95% CI u-95% CI eff.samp
## units  0.001178 0.001105 0.001246     1000
## 
##  Location effects: gr ~ mutation + anc + person + inner 
## 
##              post.mean   l-95% CI   u-95% CI eff.samp  pMCMC    
## (Intercept)  1.0879545  1.0690193  1.1060612     1000 <0.001 ***
## mutation    -0.0003071 -0.0031153  0.0026646     1086  0.882    
## anc          0.0273969 -0.0042671  0.0619536     1000  0.098 .  
## personJDH    0.0067047 -0.0213005  0.0284269     1000  0.582    
## inner       -0.0058103 -0.0092817 -0.0020537     1000  0.004 ** 
## ---
## Signif. codes:  0 '***' 0.001 '**' 0.01 '*' 0.05 '.' 0.1 ' ' 1
```

We can obtain the posterior distributions for the mean and variance of the DFE

```
mean_s<-log(2)*m1_mcmc$Sol[,"mutation"]/m1_mcmc$Sol[,"(Intercept)"]

mean(mean_s)
```

```
## [1] -0.0001941559
```

```
HPDinterval(mean_s)
```

```
##             lower       upper
## var1 -0.001979509 0.001731345
## attr(,"Probability")
## [1] 0.95
```

```
var_s<-(log(2)^2)*m1_mcmc$VCV[,"sqrt(mutation).line"]/m1_mcmc$Sol[,"(Intercept)"]^2

mean(var_s)
```

```
## [1] 1.486488e-05
```

```
HPDinterval(var_s)
```

```
##             lower        upper
## var1 7.121563e-11 5.537938e-05
## attr(,"Probability")
## [1] 0.95
```

##### 4.2.1 Estimating the DFE using the ancestor as a reference

Most studies do not have zero-mutant lines to which they can compare the fitness of lines carrying mutations. This is only valid if all between-line variance is due to mutations. If we had used the ancestor as the reference then

\[\begin{equation}
r\_w = \beta\_0+\beta\_A
\end{equation}\]

where \(\beta\_A\) is the effect of the ancestor. The mean and variance of the DFE would then be:

\[\begin{equation}
%\mu\_s = \frac{\beta\_0+\beta\_M-\beta\_0-\beta\_A}{\beta\_0+\beta\_A}ln(2)
\mu\_s = \frac{\beta\_M-\beta\_A}{\beta\_0+\beta\_A}ln(2)
\end{equation}\]

and

\[\begin{equation}
\sigma^2\_s = \frac{\sigma^2\_M}{(\beta\_0+\beta\_A)^2}ln(2)^2
\end{equation}\]

which gives

```
mean_s2<-log(2)*(m1_mcmc$Sol[,"mutation"]-m1_mcmc$Sol[,"anc"])/(m1_mcmc$Sol[,"(Intercept)"]+m1_mcmc$Sol[,"anc"])

mean(mean_s2)
```

```
## [1] -0.0170611
```

```
HPDinterval(mean_s2)
```

```
##            lower       upper
## var1 -0.03708395 0.003301582
## attr(,"Probability")
## [1] 0.95
```

```
var_s2<-(log(2)^2)*m1_mcmc$VCV[,"sqrt(mutation).line"]/(m1_mcmc$Sol[,"(Intercept)"]+m1_mcmc$Sol[,"anc"])^2

mean(var_s2)
```

```
## [1] 1.415766e-05
```

```
HPDinterval(var_s2)
```

```
##             lower        upper
## var1 6.870203e-11 5.199929e-05
## attr(,"Probability")
## [1] 0.95
```

#### 4.3 Is the ancestor effect comparable to other zero-mutant line effects?

The ancestor’s growth rate is estimated to be 0.0274 units larger than the average growth rate of the zero-mutant lines. However, the zero-mutant lines are not equivalent and vary to some degree in their growth rate - the estimated standard deviation (the square-root of the `line.1` variance) is 0.0179. Since 0.0274/0.0179 = 1.528 it is not an outlier with respect to the distribution of zero-mutant line effects. We can test this formally by fitting a random line effect for the ancestor that comes from the same distribution as the zero-mutant line effects. The fixed ancestral effect is now smaller and no longer significant:

```
m2<-lmer(gr~  mutation + anc + person + inner + (1|line) + (sqrt(mutation)-1|line) + (1|plate) + (1|plate:column) + (1|plate:row), data=df)
summary(m2)
```

```
## Linear mixed model fit by REML. t-tests use Satterthwaite's method [
## lmerModLmerTest]
## Formula: gr ~ mutation + anc + person + inner + (1 | line) + (sqrt(mutation) -  
##     1 | line) + (1 | plate) + (1 | plate:column) + (1 | plate:row)
##    Data: df
## 
## REML criterion at convergence: -9436
## 
## Scaled residuals: 
##     Min      1Q  Median      3Q     Max 
## -4.7550 -0.6015 -0.0610  0.5026  3.7585 
## 
## Random effects:
##  Groups       Name           Variance  Std.Dev. 
##  plate.column (Intercept)    4.954e-05 7.038e-03
##  plate.row    (Intercept)    1.815e-04 1.347e-02
##  line         sqrt(mutation) 1.399e-12 1.183e-06
##  line.1       (Intercept)    3.208e-04 1.791e-02
##  plate        (Intercept)    1.083e-03 3.291e-02
##  Residual                    1.172e-03 3.423e-02
## Number of obs: 2599, groups:  
## plate:column, 336; plate:row, 224; line, 188; plate, 28
## 
## Fixed effects:
##               Estimate Std. Error         df t value Pr(>|t|)    
## (Intercept)  1.088e+00  9.252e-03  2.961e+01 117.575  < 2e-16 ***
## mutation    -2.584e-04  1.373e-03  1.846e+02  -0.188  0.85090    
## anc          2.736e-02  1.869e-02  1.282e+02   1.464  0.14567    
## personJDH    6.857e-03  1.267e-02  2.603e+01   0.541  0.59293    
## inner       -5.776e-03  1.899e-03  7.404e+02  -3.041  0.00244 ** 
## ---
## Signif. codes:  0 '***' 0.001 '**' 0.01 '*' 0.05 '.' 0.1 ' ' 1
## 
## Correlation of Fixed Effects:
##           (Intr) mutatn anc    prsJDH
## mutation  -0.162                     
## anc       -0.027  0.079              
## personJDH -0.685  0.003  0.000       
## inner     -0.129 -0.001  0.008  0.000
## optimizer (nloptwrap) convergence code: 0 (OK)
## boundary (singular) fit: see help('isSingular')
```

#### 4.4 Mutation type

In the previous models all mutations are treated as being equivalent. However, we might expect synonymous mutations and small structural mutations in non-coding regions to have negligible fitness effects compared to other mutations. We can count the number of mutations per line that are not of this type (`nsyn_mutation`)

```
nsyn_mutations<-subset(mutations, (nchar(REF)!=nchar(ALT) & coding) | ALT=="<DEL>" | type%in%c("nonsynonymous", "nonsense", "frameshift"))
nnsyn_mutations<-table(factor(nsyn_mutations$line, levels=unique(df$line)))
df$nsyn_mutation<-as.numeric(nnsyn_mutations[df$line])
```

and rerun model with this as a predictor

```
m3<-lmer(gr~  nsyn_mutation + anc + person + inner + (noanc-1|line) + (sqrt(nsyn_mutation)-1|line) + (1|plate) + (1|plate:column) + (1|plate:row),data=df)
summary(m3)
```

```
## Linear mixed model fit by REML. t-tests use Satterthwaite's method [
## lmerModLmerTest]
## Formula: gr ~ nsyn_mutation + anc + person + inner + (noanc - 1 | line) +  
##     (sqrt(nsyn_mutation) - 1 | line) + (1 | plate) + (1 | plate:column) +  
##     (1 | plate:row)
##    Data: df
## 
## REML criterion at convergence: -9437.9
## 
## Scaled residuals: 
##     Min      1Q  Median      3Q     Max 
## -4.7136 -0.5992 -0.0598  0.5067  3.8460 
## 
## Random effects:
##  Groups       Name                Variance  Std.Dev.
##  plate.column (Intercept)         4.959e-05 0.007042
##  plate.row    (Intercept)         1.820e-04 0.013491
##  line         sqrt(nsyn_mutation) 9.598e-05 0.009797
##  line.1       noanc               2.641e-04 0.016252
##  plate        (Intercept)         1.083e-03 0.032910
##  Residual                         1.171e-03 0.034225
## Number of obs: 2599, groups:  
## plate:column, 336; plate:row, 224; line, 188; plate, 28
## 
## Fixed effects:
##                 Estimate Std. Error         df t value Pr(>|t|)    
## (Intercept)    1.089e+00  9.187e-03  2.877e+01 118.502  < 2e-16 ***
## nsyn_mutation -1.719e-03  2.022e-03  5.535e+01  -0.850  0.39902    
## anc            2.655e-02  5.225e-03  2.038e+03   5.082 4.08e-07 ***
## personJDH      6.837e-03  1.267e-02  2.603e+01   0.540  0.59402    
## inner         -5.819e-03  1.900e-03  7.406e+02  -3.063  0.00227 ** 
## ---
## Signif. codes:  0 '***' 0.001 '**' 0.01 '*' 0.05 '.' 0.1 ' ' 1
## 
## Correlation of Fixed Effects:
##             (Intr) nsyn_m anc    prsJDH
## nsyn_mutatn -0.112                     
## anc         -0.070  0.193              
## personJDH   -0.690  0.002  0.000       
## inner       -0.130  0.005  0.029  0.000
```

We find that even when we only look at mutations that likely have a stronger effect on fitness, mutations do not significantly influence growth rate.

#### 4.5 Significance testing of random effect variances

We use Satterthwaite’s approximate t-test for testing the significance of fixed effects (as implemented in \(\texttt{lmerTest}\)). To test the significance of the random effects we perform likelihood ratio tests implemented in the \(\texttt{anova}\) method for lmer objects (although we note that the p-values could be halved (Chernoff 1954)).

##### 4.5.1 Genetic line effects

We can omit the term `(sqrt(mutation)-1|line)` from model `m1` to test whether the variance of the DFE is non-zero

```
m4<-lmer(gr~  mutation + anc + person + inner + (noanc-1|line) + (1|plate) + (1|plate:column) + (1|plate:row),data=df)
anova(m4,m1)
```

```
## Data: df
## Models:
## m4: gr ~ mutation + anc + person + inner + (noanc - 1 | line) + (1 | plate) + (1 | plate:column) + (1 | plate:row)
## m1: gr ~ mutation + anc + person + inner + (noanc - 1 | line) + (sqrt(mutation) - 1 | line) + (1 | plate) + (1 | plate:column) + (1 | plate:row)
##    npar     AIC     BIC logLik -2*log(L) Chisq Df Pr(>Chisq)
## m4   10 -9461.9 -9403.3   4741   -9481.9                    
## m1   11 -9459.9 -9395.4   4741   -9481.9     0  1          1
```

The evidence is not strong

##### 4.5.2 Environmental line effects

We can omit the term `(noanc-1|line)` from model `m1` to test whether the variance of the EDFE is non-zero

```
m5<-lmer(gr~  mutation + anc + person + inner + (sqrt(mutation)-1|line) + (1|plate) + (1|plate:column) + (1|plate:row),data=df)
anova(m5,m1)
```

```
## Data: df
## Models:
## m5: gr ~ mutation + anc + person + inner + (sqrt(mutation) - 1 | line) + (1 | plate) + (1 | plate:column) + (1 | plate:row)
## m1: gr ~ mutation + anc + person + inner + (noanc - 1 | line) + (sqrt(mutation) - 1 | line) + (1 | plate) + (1 | plate:column) + (1 | plate:row)
##    npar     AIC     BIC logLik -2*log(L)  Chisq Df Pr(>Chisq)    
## m5   10 -9405.5 -9346.9 4712.8   -9425.5                         
## m1   11 -9459.9 -9395.4 4741.0   -9481.9 56.431  1  5.821e-14 ***
## ---
## Signif. codes:  0 '***' 0.001 '**' 0.01 '*' 0.05 '.' 0.1 ' ' 1
```

There is strong evidence that lines vary in their fitness due to things other than the mutations they’ve accumulated.

### 5 Comparison with Sane, Miranda, and Agashe (2018)

Sane, Miranda, and Agashe (2018) also used an MA procedure where lines differ by single mutations. However, rather than running many lines for a short period of time and sequencing at the end, they ran few lines for a long period and sequenced them periodically. Consequently, rather than comparing MA lines with a common unmutated ancestor, each line is compared to its ancestor prior to the accumulation of a single mutation. In total, 80 single mutations were assayed.

```
df_long_Sane<-read.csv(file.path(root, "meta/Sane_data.csv"))
```

The data frame was kindly provided by Mrudula Sane and Deepa Agashe and contains three columns `abs_gr_1`, `abs_gr_2`, `abs_gr_3` which are the three replicate growth rate estimates for each line. `cognate_anc_gr` is the estimated growth rate of that line’s immediate ancestor (presumably the average of three replicates). The `rel_gr` column is the relative growth rate as used in the paper which is the average value of the `abs_gr` variables, divided by `cognate_anc_gr` (note this is equal to \(1+(r\_i-r\_w)/r\_w\)).

We can turn this wide-format data-frame into long format with the three absolute growth rates appearing in the single column (`gr`):

```
df_Sane <- df_long_Sane[,c('mutation_id',"abs_gr_1", "abs_gr_2", "abs_gr_3","cognate_anc_gr", "transfer")] %>%
    pivot_longer(
        cols = c("abs_gr_1", "abs_gr_2", "abs_gr_3"),
        names_to = "replicate",
        values_to = "gr"
    )
```

Ideally, we would have information of the replicate estimates that formed each `cognate_anc_gr` in order to model uncertainty in \(r\_w\). It would also have been good to know the line identity of the immediate ancestor, since the line may itself have acquired a single mutation from a more distant ancestor. Since this information is not available we will analyse \(ln(2)(r\_m-r\_w)/r\_w\)

```
df_Sane=df_Sane %>% mutate(s =log(2)*(gr-cognate_anc_gr)/cognate_anc_gr)

m1_Sane<-lmer(s~(1|mutation_id),data=df_Sane)
```

```
summary(m1_Sane)
```

```
## Linear mixed model fit by REML. t-tests use Satterthwaite's method [
## lmerModLmerTest]
## Formula: s ~ (1 | mutation_id)
##    Data: df_Sane
## 
## REML criterion at convergence: -537.3
## 
## Scaled residuals: 
##     Min      1Q  Median      3Q     Max 
## -2.1895 -0.3816  0.0241  0.3808  3.6548 
## 
## Random effects:
##  Groups      Name        Variance Std.Dev.
##  mutation_id (Intercept) 0.011143 0.10556 
##  Residual                0.002506 0.05006 
## Number of obs: 240, groups:  mutation_id, 80
## 
## Fixed effects:
##             Estimate Std. Error       df t value Pr(>|t|)  
## (Intercept)  0.03127    0.01224 79.00000   2.556   0.0125 *
## ---
## Signif. codes:  0 '***' 0.001 '**' 0.01 '*' 0.05 '.' 0.1 ' ' 1
```

The mean of the DFE (the intercept) exceeds zero by some margin and the variance of the DFE is also substantial and highly significant:

```
anova(m1_Sane, lm(s~1,data=df_Sane))
```

```
## Data: df_Sane
## Models:
## lm(s ~ 1, data = df_Sane): s ~ 1
## m1_Sane: s ~ (1 | mutation_id)
##                           npar     AIC     BIC logLik -2*log(L)  Chisq Df
## lm(s ~ 1, data = df_Sane)    2 -348.13 -341.17 176.07   -352.13          
## m1_Sane                      3 -538.25 -527.81 272.12   -544.25 192.12  1
##                           Pr(>Chisq)    
## lm(s ~ 1, data = df_Sane)               
## m1_Sane                    < 2.2e-16 ***
## ---
## Signif. codes:  0 '***' 0.001 '**' 0.01 '*' 0.05 '.' 0.1 ' ' 1
```

Part of the between-line variance will be driven by the fact that the same `cognate_anc_gr` value is used for each of the three replicate growth rates of a mutation. However, if we analyse the replicate growth rates of each line without normalising by the ancestors growth rate and generation time we still see substantial between-line variance.

```
m2_Sane<-lmer(gr~(1|mutation_id),data=df_Sane)
```

```
summary(m2_Sane)
```

```
## Linear mixed model fit by REML. t-tests use Satterthwaite's method [
## lmerModLmerTest]
## Formula: gr ~ (1 | mutation_id)
##    Data: df_Sane
## 
## REML criterion at convergence: -286.3
## 
## Scaled residuals: 
##     Min      1Q  Median      3Q     Max 
## -2.3529 -0.4401  0.0288  0.3970  3.5153 
## 
## Random effects:
##  Groups      Name        Variance Std.Dev.
##  mutation_id (Intercept) 0.022953 0.15150 
##  Residual                0.008253 0.09085 
## Number of obs: 240, groups:  mutation_id, 80
## 
## Fixed effects:
##             Estimate Std. Error       df t value Pr(>|t|)    
## (Intercept)  1.33697    0.01792 79.00000   74.59   <2e-16 ***
## ---
## Signif. codes:  0 '***' 0.001 '**' 0.01 '*' 0.05 '.' 0.1 ' ' 1
```

We can also fit the model `m1_Sane` using Markov chain Monte Carlo:

```
m1_Sane_mcmc<-MCMCglmm(s~1, 
                       random=~mutation_id, 
                       data=df_Sane, 
                       prior=list(R=IW(1,0.002), G=F(1,1000)), 
                       longer=10,
                      verbose=FALSE) 

summary(m1_Sane_mcmc)
```

```
## 
##  Iterations = 30001:129901
##  Thinning interval  = 100
##  Sample size  = 1000 
## 
##  DIC: -679.3643 
## 
##  G-structure:  ~mutation_id
## 
##             post.mean l-95% CI u-95% CI eff.samp
## mutation_id   0.01159 0.008193  0.01611     1000
## 
##  R-structure:  ~units
## 
##       post.mean l-95% CI u-95% CI eff.samp
## units  0.002543    0.002 0.003081     1000
## 
##  Location effects: s ~ 1 
## 
##             post.mean l-95% CI u-95% CI eff.samp pMCMC  
## (Intercept)  0.031139 0.005869 0.056394    755.2 0.024 *
## ---
## Signif. codes:  0 '***' 0.001 '**' 0.01 '*' 0.05 '.' 0.1 ' ' 1
```

Blazanin, Michael. 2024. “gcplyr: an R package for microbial growth curve data analysis.” *BMC Bioinformatics* 25 (232).

Chernoff, Herman. 1954. “On the Distribution of the Likelihood Ratio.” *The Annals of Mathematical Statistics*, 573–78.

Chevin, Luis-Miguel. 2011. “On Measuring Selection in Experimental Evolution.” *Biology Letters* 7 (2): 210–13.

Fink, Justus Wilhelm, and Michael Manhart. 2024. “Quantifying Microbial Fitness in High-Throughput Experiments.” *bioRxiv*, 2024–08.

Sane, Mrudula, Joshua John Miranda, and Deepa Agashe. 2018. “Antagonistic Pleiotropy for Carbon Use Is Rare in New Mutations.” *Evolution* 72 (10): 2202–13.
