## Supplementary material for "Fitness effects of new mutations are small and heavily confounded with non-genetic sources of variation in Escherichia coli": Mutation Calling: Mutation_Calling.html


### Mutation Calling

- 1 Mapping Reads
- 2 Calling Mutations
- 3 Annotating Mutations
- 4 Filtering Mutations
  - 4.1 Ancestral mutations
  - 4.2 Variant calls with low Phred-scaled score (QUAL\(<20\))
  - 4.3 Possible mutations
  - 4.4 BPS with less than 95% support
  - 4.5 INDELS with less than 95% support
  - 4.6 BPS with fewer than 30 supporting reads
  - 4.7 INDELS with fewer than 30 supporting reads
  - 4.8 rRNA and insertion sequence
  - 4.9 Large structural variants
  - 4.10 Real mutations
- 5 False-negative rate for mutation calling
  - 5.1 Simulate mutations
  - 5.2 Call simulated mutations

In this workbook we detail the workflow for mapping reads and calling, validating and annotating variants in 194 *E. coli* mutation accumulation lines and two replicates of their common ancestor. The raw reads can be found on NCBI’s SRA (see BioProject ID PRJNA1466858). A root directory needs to be specified (`~/Work/Ecoli_MA`) and within that directory a subdirectory called raw should exist. The fasta files are to large to be hosted on GitHib but can be downloaded from NCBI’s Sequence Read Archive under BioProject ID PRJNA1466858. They should be placed in the raw directory with the fasta files for each library contained within its own subdirectory. The subdirectory should be named after the sample. The sample name is the leading text of the SRA Name field prior to the sequence and the MA line is listed in the SRA Title field (e.g the first SRA is sample 270-94 corresponding to MA Line 26). Details about the samples can be found in meta/sample\_submission.csv.

### 1 Mapping Reads

The reads are mapped to the specified reference genome (NCBI RefSeq accession NC\_000913.3, assembly GCF\_000005845.2 (ASM584v2)) using bwa (Li 2013). bam files are stored in a subdirectory of root called bam. If the bam file for a sample exists mapping will be skipped. The bash script map.sh (in the scripts subdirectory of root) is shown below.

```
#!/usr/bin/env bash
set -euo pipefail

# ------------- USER SETTINGS ---------------------------------
bwa_threads=8
export ref=~/Work/Ecoli_MA/ref/GCF_000005845.2_ASM584v2_genomic.fna # reference genome
root=~/Work/Ecoli_MA                                                # project root
# -------------------------------------------------------------

if [[ ! -f ${ref}.bwt ]]; then bwa index  "$ref"; fi
if [[ ! -f ${ref}.fai ]]; then samtools faidx "$ref"; fi
# create bwa and fai index files for the reference genome if they do not exist

mkdir -p "$root"/bam  # directory for storing bam files  

raw="$root"/raw  # location of fastq files (separate directories per sample)

echo -e "\n=== ALIGNMENT ==="

for dir in "$raw"/*/ ; do                     # every sample folder
    [[ -d $dir ]] || continue                 # check dir is a directory

    sample=${dir#"$raw"/} ; sample=${sample%/}  # get sample name
    bam=$root/bam/${sample}.bam                 # output bam file name
    [[ -f $bam ]] && { echo "SKIPPED: $sample already aligned" ; continue ; } # skips sample if bam file already exists

    r1=$(find "$dir" -maxdepth 1 -name '*_R1*.fastq.gz' -print -quit)
    r2=$(find "$dir" -maxdepth 1 -name '*_R2*.fastq.gz' -print -quit)
    # get file names for the paired ends

    if [[ -z "$r1" || -z "$r2" ]]; then
        echo "PROBLEM: Missing R1 or R2 file in $dir – skipped" >&2
        continue
    fi

    echo "ALLIGNING  $sample"

    bwa mem -t"$bwa_threads" "$ref"  "$r1" "$r2" \
      | samtools sort -n  -@4 -o - -          \
      | samtools fixmate   -m - -             \
      | samtools sort      -@4 -o - -         \
      | samtools markdup   -@4 - "$bam"    

    # aligns to reference genome
    # sorts output by read name
    # corrects mate-pair info
    # resorts output by location
    # mark duplicate reads     

    samtools index "$bam"   # create index file for the final BAM file


done

echo -e "\n Mapping finished OK"
```

```
system(file.path(root, "scripts/map.sh"))
```

### 2 Calling Mutations

PCR duplicates, secondary reads and non-mapped reads were removed using samtools and variants called using bcftools (Danecek et al. 2021). The bash script call.sh (in the scripts subdirectory of root) is shown below.

```
#!/usr/bin/env bash
set -euo pipefail

# ------------- USER SETTINGS ---------------------------------
threads=32                               # number of cores to be used                   
export ref=~/Work/Ecoli_MA/ref/GCF_000005845.2_ASM584v2_genomic.fna  # reference genome
root=~/Work/Ecoli_MA                    # project root
# -------------------------------------------------------------

mkdir -p "$root"/vcf
# directory for writing vcf files


echo -e "\n=== VARIANT CALLING ==="
parallel -j2 '
    bam={}                                                    # path to bam file
    sample=$(basename ${bam%.bam})                            # generates sample name from bam file name
    rawvcf='$root'/vcf/${sample}.raw.vcf.gz                   # path to vcf output file
    filtered_bam='$root'/filtered_bam/${sample}.filtered.bam           # path to filtered bam file

    [[ -f $rawvcf ]] && { echo "SKIPPED  $sample raw VCF exists" ; exit; }

    samtools view -F 0xF04 -b "$bam" -o "$filtered_bam"
    samtools index "$filtered_bam"

    # removes unmapped, QC-failed and duplicate reads + secondary/supplementary alignments

    bcftools mpileup  --threads 4 -f '"$ref"' \
        -E  \
        --min-MQ 30 --min-BQ 20 "$filtered_bam" |   
    bcftools call   --threads 4 --ploidy 1 -m -Oz -o "$rawvcf"

    # summarise the bases at aligned reads only using reads with mapping quality >30
    # and at bases with base quality >20. Then call variants
    
    bcftools index  "$rawvcf"  # index the vcf files
' ::: "$root"/bam/*.bam

echo -e "\n=== FILTERING ==="

parallel -j"$threads" '
    
    invcf={}                               # path to vcf files
    
    sample=$(basename "$invcf" .raw.vcf.gz)  # get sample name from vcf file name

    outvcf='$root'/vcf/${sample}.flt.vcf.gz  # path to filtered vcf output file

    [[ -f $outvcf ]] && { echo "SKIPPED  $sample filtered VCF exists" ; exit ; }

    bcftools filter -e "QUAL<20" -S . "$invcf" | 
    bcftools view   -v snps,indels -Ou |
    bcftools norm  -m -any -f "$ref" -Oz -o "$outvcf"

    # flags sites with a QUAL score less than 20
    # filter out variants that are not snps or indels
    # standardises variant format
    # index the filtered vcf files
    # standardises variant format

    bcftools index "$outvcf"  # index the fitered vcf files

' ::: "$root"/vcf/*.raw.vcf.gz

echo -e "\n Variant calling finished OK"
```

```
system(file.path(root, "scripts/call.sh"))
```

### 3 Annotating Mutations

We use an R pipeline to filter and annotate variants. Variants at sites with a Phred-scaled score (QUAL) \(<20\) are excluded and the remainder are considered possible mutations. These possible mutations and associated meta-data are written to pos\_mutations.csv in a subdirectory of root called meta. Possible mutations with at least 30 reads supporting the alternate allele and less than 5% of reads supporting the reference allele were considered real mutations without the need for checking. Possible mutations not meeting this strict filter were visualised in the Integrative Genomics Viewer (Robinson et al. 2011). The file manual\_check.csv in the subdirectory meta states which of the possible mutations were checked and whether they were validated or not. Using this information a set of real validated mutations are written to real\_mutations.csv in the meta subdirectory.

We can start by organising the necessary objects for annotation

```
library(data.table)
library(VariantAnnotation)  
library(GenomicFeatures) 
library(rtracklayer)      
library(txdbmaker)
library(GenomeInfoDb)

sample_line<-read.csv(file.path(root, "meta/sample_submission.csv"))
# meta-data for samples 

gff_file <- file.path(root, "ref/GCF_000005845.2_ASM584v2_genomic.gff")
fna_file <- file.path(root, "ref/GCF_000005845.2_ASM584v2_genomic.fna")    

# path to gff genome annotation file and FASTA nucleotide sequence for the reference

txdb <- makeTxDbFromGFF(gff_file, format = "gff")
# create database of genomic features

gff <- import(gff_file)
# import the gff file

cds_gr <- cds(txdb,  columns = c("gene_id", "tx_name"))
# extract coding sequence regions

pseudo_gr <- gff[gff$type=="pseudogene"]
rrna_gr <- gff[gff$type=="rRNA"]
is_gr <- gff[grepl("insertion sequence", gff$mobile_element_type)]
gene_gr <- gff[gff$type=="gene"]

# filter gff files for pseudogenes, rrna, insertion sequences and genes

fa <- FaFile(fna_file)  
# creates a FaFile object for efficiently accessing the FASTA file
```

Then we work through each vcf file, extract the relevant information and annotate the mutations.

```
vcf_files<-list.files(file.path(root, "vcf"))
vcf_files<-vcf_files[grep(".flt.vcf.gz$", vcf_files)]
# get a list filtered vcf file names 

sample<-unlist(lapply(strsplit(vcf_files, "_|\\."), function(x){x[2]}))
sample<-gsub("391-Ancestor", "391-Ancestor_1", sample)
sample<-gsub("392-Ancestor", "392-Ancestor_2", sample)
# get the sample names from the vcf file names. 

line<-sample_line$Sample.name[match(unlist(lapply(strsplit(sample, "-"), function(x){x[1]})), sample_line$Sample.number)]
# get the MA line names associated with each sample. 

vcf_dt<-c()

for(i in 1:length(vcf_files)){

    file.i<-file.path(root, "vcf", vcf_files[i])
    vcf <- readVcf(file.i)
    # read in the filtered vcf file

    vr  <- granges(vcf)                                                 # get the coordinates of the variants

    is_cds<-countOverlaps(vr, cds_gr)>0
  is_pseudo<-countOverlaps(vr, pseudo_gr)>0
  is_rrna<-countOverlaps(vr, rrna_gr)>0
  is_is<-countOverlaps(vr, is_gr)>0

  # evaluate whether the variants overlap coding sequence, 
  # pseudogenes, rrna genes and insertion sequences.

    coding <- predictCoding(vcf, txdb, seqSource = fa)
  # gets coding mutation types (synonymous, non-synonymous, )

    mini_vcf<-fread(file.i, sep = "\t", header = TRUE, skip = "#CHROM")
  # read vcf into a data.table

    n_ref<-as.vector(sapply(mini_vcf$INFO, function(x){sum(as.numeric(strsplit(strsplit(x, "DP4=|;MQ=")[[1]][2], ",")[[1]])[1:2])}))
    n_alt<-as.vector(sapply(mini_vcf$INFO, function(x){sum(as.numeric(strsplit(strsplit(x, "DP4=|;MQ=")[[1]][2], ",")[[1]])[3:4])}))
  # get number of reads in support of the reference and alternate allele

    is_nonsyn<-gene<-locus_tag<-rep(NA, length(names(vcf)))
    # empty vector for writing mutation type and gene name for mutations in coding sequence

    is_nonsyn[match(names(coding), names(vcf))]<-as.character(coding$CONSEQUENCE)
  # writes mutation type for mutations in coding sequence

  hits    <- findOverlaps(vr, gene_gr)
  # finds overlap between variant coordinates and genes

  gene[queryHits(hits)] <- unlist(gene_gr$Name[subjectHits(hits)])
  locus_tag[queryHits(hits)] <- unlist(gene_gr$locus_tag[subjectHits(hits)])
  # writes gene name for mutations falling in genes 

    vcf_dt <- rbind(vcf_dt, cbind(sample[i], line[i], is_cds, is_nonsyn, is_pseudo, is_rrna, is_is, gene, locus_tag, mini_vcf$QUAL, n_ref, n_alt, mini_vcf[,c(2,4,5)]))

  # consolidate information into a data.frame 
}

colnames(vcf_dt)[1:12]<-c("sample", "line", "coding", "type", "pseudo", "rrna", "is", "gene", "locus_tag", "QUAL", "n_ref", "n_alt")
```

### 4 Filtering Mutations

`vcf_dt` contains 2125 variant calls that have not been filtered in any way. We first filter out the two variants that are present in the ancestor and therefore all MA lines. We then filter out sites of low quality to give us a set of possible mutations. These possible mutations are split in to a high confidence set (more than 30 reads and more than 95% of reads supporting the variant) and a low confidence set. The low confidence set are manually validated, and information is given as to why we think they are valid or not.

#### 4.1 Ancestral mutations

Every line had an identical variant called at positions 4296380 and 4381253. These are mutations that have occurred in our ancestral line before MA commenced and so we remove them:

```
orig_mut<-c(4296380, 4381253)

vcf_dt<-subset(vcf_dt, !POS%in%orig_mut)
```

#### 4.2 Variant calls with low Phred-scaled score (QUAL\(<20\))

For the remaining 1733 variant calls, we can plot the proportion of reads that support the alternate allele against the number of reads that support the alternate allele. In black we have INDELS and in red we have bps. Triangles are mutations in rRNA genes or insertion sequences, and circles are the remainder.

```
vcf_dt$prop<-vcf_dt$n_alt/(vcf_dt$n_ref+vcf_dt$n_alt)
plot(jitter(prop, amount=0.01)~jitter(n_alt,5), col=1+as.numeric(nchar(ALT)==nchar(REF)), cex=0.5, ylab=c("Proportion alternate"),  pch=c(1,17)[1+as.numeric(rrna | is)], xlab=c("Number alternate"), data=vcf_dt)
```

Note that most variant calls only have a single read in support of the alternate allele:

```
table(vcf_dt$n_alt==1)
```

```
## 
## FALSE  TRUE 
##   263  1470
```

Of those that only have a single read in support of the alternate allele, most are either in rRNA or insertion sequences

```
table(subset(vcf_dt, n_alt==1)$rrna | subset(vcf_dt, n_alt==1)$is)
```

```
## 
## FALSE  TRUE 
##   178  1292
```

If we exclude sites with low quality, most of these sites disappear and we are left with 255 possible mutations:

```
vcf_dt_qual<-subset(vcf_dt, QUAL>20)
```

#### 4.3 Possible mutations

For variants that pass the quality filter, we can see that bps generally have close to 100% of reads supporting the alternate, whereas for INDELS the proportion is generally less.

```
plot(jitter(prop, amount=0.01)~jitter(n_alt,2), col=1+as.numeric(nchar(ALT)==nchar(REF)), cex=1, ylab=c("Proportion alternate"),  pch=c(1,17)[1+as.numeric(rrna | is)], xlab=c("Number alternate"), data=vcf_dt_qual)
abline(v=30, lty=2)
abline(h=0.95, lty=2)
```

with at least 30 reads supporting the alternate allele and less than 5% of reads supporting the reference allele

All mutations with with at least 30 reads in support of the alternate allele and more than 95% of reads in support of the alternate allele (i.e. the upper right-hand quadrant formed by the dotted lines) were considered unambiguously real. The remainder were checked in the Integrative Genomics Viewer (Robinson et al. 2011). The `check` column of manual\_check.csv indicates whether the mutation was checked (not `NA`) and whether it was deemed OK (`OK`) or not (`NO`).

```
manual_check<-read.csv(file.path(root, "meta/manual_check.csv"))

pos_in_man<-match(paste(vcf_dt_qual$sample,vcf_dt_qual$POS), paste(manual_check$sample,manual_check$POS))

vcf_dt_qual$check<-manual_check$check[pos_in_man]
```

Note that 271-96 1196220, which met the strict filter, was also checked as it was in *icd* and had exactly 30 reads in support (see below)

#### 4.4 BPS with less than 95% support

We can look at the 6 bps with 95% of read support or less:

```
subset(vcf_dt_qual[,-c(2:5)], prop<=0.95 & nchar(ALT)==nchar(REF))
```

```
##     sample   rrna     is   gene locus_tag    QUAL n_ref n_alt     POS    REF
##     <char> <lgcl> <lgcl> <char>    <char>   <num> <num> <num>   <int> <char>
## 1:  202-10   TRUE  FALSE   rrlH     b0204 228.352     1    19  226642      T
## 2:  230-42  FALSE  FALSE   <NA>      <NA> 117.077    13    67  960164      T
## 3: 279-107  FALSE  FALSE   ynfO     b4533 228.236     5    50 1636981      C
## 4: 279-107  FALSE  FALSE   letB     b1834 228.234     4    43 1918129      T
## 5: 287-116  FALSE  FALSE   caiC     b0037 228.270     7    83   37246      A
## 6: 287-116   TRUE  FALSE   rrsH     b0201 123.251     1    13  223974      G
##       ALT      prop  check
##    <char>     <num> <char>
## 1:      G 0.9500000     OK
## 2:      G 0.8375000     NO
## 3:      T 0.9090909     OK
## 4:      A 0.9148936     OK
## 5:      C 0.9222222     OK
## 6:      T 0.9285714     NO
```

Two looked like mapping error (one is in a rRNA gene) although one of those deemed as an error (230-42 960164 T->G) could have been real if there was a lot of contamination - hard to verify since the line didn’t have other mutations. Four looked OK, but 202-10 226642 T->G is in an rRNA gene and so we removed it

```
vcf_dt_qual$check[which(vcf_dt_qual$sample=="202-10" & vcf_dt_qual$POS=="226642")]<-"NO"
```

The two 279-107 mutations looked OK and there was perhaps some contamination as all mutations had some reads supporting the reference

```
subset(vcf_dt_qual[,-c(2:5)], sample=="279-107")
```

```
##     sample   rrna     is   gene locus_tag    QUAL n_ref n_alt     POS    REF
##     <char> <lgcl> <lgcl> <char>    <char>   <num> <num> <num>   <int> <char>
## 1: 279-107  FALSE  FALSE   ynfO     b4533 228.236     5    50 1636981      C
## 2: 279-107  FALSE  FALSE   letB     b1834 228.234     4    43 1918129      T
## 3: 279-107  FALSE  FALSE   cirA     b2155 228.386     1    50 2245042      C
##       ALT      prop  check
##    <char>     <num> <char>
## 1:      T 0.9090909     OK
## 2:      A 0.9148936     OK
## 3:      A 0.9803922     OK
```

287-116 37246 looked OK - perhaps some contamination also.

#### 4.5 INDELS with less than 95% support

28 INDELS have less than 95% of read support. None occurred in rRNA genes or insertion sequences.

```
subset(vcf_dt_qual[,-c(2:5)], prop<=0.95 & nchar(ALT)!=nchar(REF))
```

```
##      sample   rrna     is   gene locus_tag     QUAL n_ref n_alt     POS
##      <char> <lgcl> <lgcl> <char>    <char>    <num> <num> <num>   <int>
##  1:  204-12  FALSE  FALSE   <NA>      <NA> 159.9400    36    38 1198453
##  2:  222-30  FALSE  FALSE   <NA>      <NA> 171.1400     7    28  380012
##  3:  232-44  FALSE  FALSE   <NA>      <NA> 216.0890    40    61 1198453
##  4:  242-57  FALSE  FALSE   fimZ     b0535 228.3190     8    65  564099
##  5:  253-74  FALSE  FALSE   <NA>      <NA> 143.3650     2    31 4166465
##  6:  255-76  FALSE  FALSE   dgcE     b2067 228.3750     5    76 2144783
##  7:  262-84  FALSE  FALSE   <NA>      <NA> 131.7670    28    22 1198436
##  8:  266-90  FALSE  FALSE   tdcG     b4471 228.3120     5    41 3258688
##  9:  272-97  FALSE  FALSE   <NA>      <NA> 120.6340    43    26 2765411
## 10: 298-128  FALSE  FALSE   <NA>      <NA> 221.1420    30    50 2765411
## 11: 310-142  FALSE  FALSE   ydeI     b1536 228.3530     4    58 1624349
## 12: 310-142  FALSE  FALSE   yoaA     b1808 228.3440     8    65 1892648
## 13: 310-142  FALSE  FALSE   <NA>      <NA> 228.3710     5    83 1986327
## 14: 310-142  FALSE  FALSE   <NA>      <NA> 145.2740     7    29 2185868
## 15: 310-142  FALSE  FALSE   leuV     b4368 228.3000     9    60 4606086
## 16: 312-145  FALSE  FALSE   <NA>      <NA> 219.1410    27    48 1198453
## 17: 333-167  FALSE  FALSE   gabD     b2661 228.3970     4    69 2791836
## 18: 344-178  FALSE  FALSE   mlaD     b3193 228.3640     7    87 3338444
## 19: 346-180  FALSE  FALSE   <NA>      <NA> 228.3400     6    75 2151059
## 20: 347-181  FALSE  FALSE   lsrG     b1518 228.3530     6    64 1607059
## 21: 369-204  FALSE  FALSE   <NA>      <NA> 228.3890     6    93 3941551
## 22: 374-210  FALSE  FALSE   wzzB     b2027 228.3210    15    79 2098233
## 23: 376-212  FALSE  FALSE   waaZ     b3624 228.3510     8    90 3799382
## 24: 378-214  FALSE  FALSE   <NA>      <NA> 228.2780    12    70 1212080
## 25:  384-33  FALSE  FALSE   ybhL     b0786 210.1360    28    49  820184
## 26: 389-209  FALSE  FALSE   <NA>      <NA> 195.9560    36    36 2560977
## 27: 389-209  FALSE  FALSE   <NA>      <NA>  48.9103    50    14 2560999
## 28:  390-38  FALSE  FALSE   <NA>      <NA> 109.8210    34    32 2765411
##      sample   rrna     is   gene locus_tag     QUAL n_ref n_alt     POS
##      <char> <lgcl> <lgcl> <char>    <char>    <num> <num> <num>   <int>
##                                       REF       ALT      prop  check
##                                    <char>    <char>     <num> <char>
##  1:                                     A AATGAAATG 0.5135135     NO
##  2:                                    AG         A 0.8000000     OK
##  3:                             AATGAAATG         A 0.6039604     NO
##  4:                                    GA         G 0.8904110     OK
##  5:                                    GA         G 0.9393939     OK
##  6:                                     C        CA 0.9382716     OK
##  7:                                     A AAATGAAAT 0.4400000     NO
##  8:                                    GC         G 0.8913043     OK
##  9:                                     C CGCACTATG 0.3768116     NO
## 10:                             CGCACTATG         C 0.6250000     NO
## 11:                                    AT         A 0.9354839     OK
## 12:                                  TACC         T 0.8904110     OK
## 13:                                    GA         G 0.9431818     OK
## 14:                                     T        TG 0.8055556     OK
## 15:                                    AG         A 0.8695652     OK
## 16:                             AATGAAATG         A 0.6400000     NO
## 17:                               TGCGCTG         T 0.9452055     OK
## 18:                                    AT         A 0.9255319     OK
## 19:                                    GA         G 0.9259259     OK
## 20:                                    GT         G 0.9142857     OK
## 21:                                     C        CA 0.9393939     OK
## 22:                            TAAATCAATC         T 0.8404255     OK
## 23:                                    AT         A 0.9183673     OK
## 24:                                     A        AC 0.8536585     OK
## 25: TCCAGTATATTCATTGTCTATACCGCTGCTTCTATCG         T 0.6363636     OK
## 26:              TGGTGGGTCAAAAGTTGCCGTTAA         T 0.5000000     OK
## 27:                                     A        AG 0.2187500     NO
## 28:                                     C CGCACTATG 0.4848485     NO
##                                       REF       ALT      prop  check
##                                    <char>    <char>     <num> <char>
```

The mutations

204-12 1198453 A->AATGAAATG

262-84 1198436 A->AAATGAAAT

312-145 1198453 AATGAAATG->A

232-44 1198453 AATGAAATG->A

were all deemed mapping errors as they were in same region across lines. Similarly,

272-97 2765411 C->CGCACTATG

298-128 2765411 CGCACTATG->C

390-38 2765411 C->CGCACTATG

were deemed mapping errors for the same reason.

389-209 2560977 TGGTGGGTCAAAAGTTGCCGTTAA->T was deemed real but the mutation 389-209 2560999 A->AG unlikely to be real - only one mutation happened in this region.

The mutations

310-142 1624349 AT->A

310-142 1892648 TACC->T

310-142 1986327 GA->G

310-142 4606086 AG->A

310-142 2185868 T->TG

were all deemed real, but sample 310-142 has 17 additional mutations, one in the mismatch repair gene *mutS*. There’s no sign of contamination - it seems like these mutations occurred after the conclusion of mutation accumulation but during the growth stage prior to sequencing.

The remaining mutations were all deemed correct. In most cases the INDEL fell in a repetitive region and the reads supporting the reference did not span sufficient repeats to call the mutation.

#### 4.6 BPS with fewer than 30 supporting reads

We can also check the 24 bps with fewer than 30 reads in support of the alternate, yet the proportion exceeded 95%:

```
subset(vcf_dt_qual[,-c(2:5)], n_alt<30 & prop>0.95 & nchar(ALT)==nchar(REF))
```

```
##      sample   rrna     is   gene locus_tag     QUAL n_ref n_alt     POS    REF
##      <char> <lgcl> <lgcl> <char>    <char>    <num> <num> <num>   <int> <char>
##  1:  211-19   TRUE  FALSE   rrsD     b3278  27.4222     0     2 3427350      T
##  2:  215-23  FALSE   TRUE  insI2     b1404 108.4150     0     6 1470138      C
##  3:  224-41  FALSE  FALSE   <NA>      <NA> 225.4170     0    18 2521360      A
##  4:  237-50  FALSE  FALSE   <NA>      <NA>  37.4152     0     2 2878065      T
##  5:  238-53  FALSE   TRUE insH10     b3218  37.4152     0     2 3366187      G
##  6:  250-71   TRUE  FALSE   rrsA     b3851 108.4150     0     6 4036756      C
##  7:  263-85  FALSE  FALSE   <NA>      <NA> 112.4150     0     6 3946620      A
##  8:  271-96  FALSE  FALSE    icd     b1136 225.4170     0    26 1196232      C
##  9:  271-96  FALSE  FALSE    icd     b1136 214.4170     0    17 1196245      T
## 10:  271-96  FALSE  FALSE    icd     b1136 213.4170     0    15 1196247      A
## 11:  271-96  FALSE  FALSE    icd     b1136 152.4160     0    12 1196277      C
## 12:  271-96  FALSE  FALSE    icd     b1136 132.4160     0     8 1196280      G
## 13:  271-96  FALSE  FALSE    icd     b1136 132.4160     0     8 1196283      A
## 14:  271-96  FALSE  FALSE    icd     b1136 129.4160     0     8 1196292      C
## 15:  271-96  FALSE  FALSE    icd     b1136  93.4151     0     5 1196304      G
## 16:  271-96  FALSE  FALSE    icd     b1136  62.4147     0     3 1196316      T
## 17: 294-124  FALSE  FALSE   yafN     b0232  30.4183     0     1  252040      C
## 18: 300-130  FALSE   TRUE  insH7     b2030  27.4222     0     2 2102455      A
## 19: 314-147   TRUE  FALSE   rrlC     b3758  27.4222     0     2 3944577      G
## 20: 328-162  FALSE  FALSE   <NA>      <NA> 225.4170     0    23 3943420      G
## 21: 338-172  FALSE  FALSE   <NA>      <NA> 178.4160     0     8 2818046      A
## 22: 338-172  FALSE  FALSE   <NA>      <NA> 178.4160     0     8 2818049      T
## 23: 339-173   TRUE  FALSE   rrlA     b3854 129.4160     0     9 4038402      T
## 24: 358-192  FALSE   TRUE  insL2     b0582  37.4152     0     2  608506      A
##      sample   rrna     is   gene locus_tag     QUAL n_ref n_alt     POS    REF
##      <char> <lgcl> <lgcl> <char>    <char>    <num> <num> <num>   <int> <char>
##        ALT  prop  check
##     <char> <num> <char>
##  1:      G     1     NO
##  2:      G     1     NO
##  3:      G     1     OK
##  4:      C     1     NO
##  5:      T     1     NO
##  6:      A     1     NO
##  7:      G     1     OK
##  8:      T     1     NO
##  9:      C     1     NO
## 10:      G     1     NO
## 11:      T     1     NO
## 12:      C     1     NO
## 13:      G     1     NO
## 14:      T     1     NO
## 15:      A     1     NO
## 16:      A     1     NO
## 17:      G     1     NO
## 18:      C     1     NO
## 19:      T     1     NO
## 20:      A     1     OK
## 21:      G     1     OK
## 22:      A     1     OK
## 23:      G     1     OK
## 24:      C     1     NO
##        ALT  prop  check
##     <char> <num> <char>
```

Four are in an rRNA gene and four are in an insertion sequence. Nine mutations are in the same line in the *icd* gene and were considered mapping errors (see below). Of the remaining six, four looked real (minimum read support is 6) and two did not (read supports of one and two)

#### 4.7 INDELS with fewer than 30 supporting reads

Three INDELS had fewer than 30 supporting reads but a proportion in excess of 0.95.

```
subset(vcf_dt_qual[,-c(2:5)], n_alt<30 & prop>0.95 & nchar(ALT)!=nchar(REF))
```

```
##     sample   rrna     is   gene locus_tag     QUAL n_ref n_alt     POS    REF
##     <char> <lgcl> <lgcl> <char>    <char>    <num> <num> <num>   <int> <char>
## 1:  252-73  FALSE  FALSE   tyrR     b1323  40.4148     0     2 1386909      G
## 2:  263-85  FALSE  FALSE   <NA>      <NA> 120.4150     0     6 3946632      A
## 3: 365-200   TRUE  FALSE   rrlB     b3970 111.4150     0     8 4168938     GA
##       ALT  prop  check
##    <char> <num> <char>
## 1:     GC     1     NO
## 2:     AT     1     OK
## 3:      G     1     NO
```

252-73 1386909 G->GC was deemed not real (only 2 reads with support) as it was adjoining a big region with no reads mapping - which was identified as a big deletion by Parliment.

365-200 4168938 GA->G was deemed not real (6 reads with support)

263-85 3946632 A->AT was deemed probably real (8 reads with support) and mapping was likely difficult due to a bps mutation at 3946620.

#### 4.8 rRNA and insertion sequence

In addition there is one bps mutation in an rRNA gene and an insertion sequence. Both have good support and are deemed real.

```
subset(vcf_dt_qual[,-c(2:5)], n_alt>30 & prop>0.95 & (rrna | is))
```

```
##     sample   rrna     is   gene locus_tag    QUAL n_ref n_alt     POS    REF
##     <char> <lgcl> <lgcl> <char>    <char>   <num> <num> <num>   <int> <char>
## 1:  262-84  FALSE   TRUE   <NA>      <NA> 225.417     0    48 1294708      C
## 2: 314-147   TRUE  FALSE   rrlC     b3758 225.421     0    60 3946577      C
##       ALT  prop  check
##    <char> <num> <char>
## 1:      G     1     OK
## 2:      G     1     OK
```

#### 4.9 Large structural variants

Three large deletions were also found using Parliament2 (Zarate et al. 2021): a 321bp deletion in sample 252-73 (1386588-1386909), a 15112bp deletion in sample 271-96 (1196300-1211412) and a 456bp deletion in sample 338-172 (2817640-2818096). All were checked in IGV. Note that the deletion in sample 271-96 spans *icd* and the false mutations called were around the ends of this deletion. The deletion coincides with the cryptic prophage e14, which has previously been shown to excise on induction of the SOS pathway (Brody, Greener, and Hill 1985).

#### 4.10 Real mutations

We only retain those mutations that passed our criteria, and excluded the mutations in the hyper-mutator that arose after mutation accumulation

```
mutations_real<-subset(vcf_dt_qual, is.na(check) | check=="OK" & !(sample=="310-142" & POS%in%c(1624349,1892648,1986327,4606086,2185868)))
```

Note that if we had simply discarded all mutations in insertion sequence, rRNA genes and the gene *icd* we would have had a reasonable agreement:

```
with(vcf_dt_qual, table(problem=(is | rrna | gene=="icd"), pass=(is.na(check) | check=="OK")))
```

```
##        pass
## problem FALSE TRUE
##   FALSE     2  166
##   TRUE     20    3
```

We add the large deletions manually to the list of real mutations:

```
tmp<-matrix(NA, 3, ncol(mutations_real))
colnames(tmp)<-colnames(mutations_real)
tmp[,"sample"]<-c("252-73", "271-96", "338-172")
tmp[,"line"]<-c(115,42,215)
tmp[,"POS"]<-c(1386588, 1196300, 2817640)
tmp[,"ALT"]<-rep("<DEL>",3)
mutations_real<-rbind(mutations_real, tmp)
```

and the save the files

```
write.csv(vcf_dt_qual, file.path(root, "meta/pos_mutations.csv"))
write.csv(mutations_real, file.path(root, "meta/real_mutations.csv"))
```

### 5 False-negative rate for mutation calling

To determine how many mutations our pipeline may have missed, we took the reference sequence and randomly changed a base, added a base and deleted a base. We then mapped reads from one of our lines to this modified reference and called variants. We repeated this process for each of the 196 libraries and assessed how often the simulated mutations were called.

#### 5.1 Simulate mutations

```
library(stringi)

bases<-c("A", "C", "T", "G")

ref_fna<-scan(file.path(root, "ref/GCF_000005845.2_ASM584v2_genomic.fna"), what="character", sep="\n")
# read in original reference genome

header<-ref_fna[1]
ref_fna<-paste(ref_fna[-1], collapse="")

genome_size<-nchar(ref_fna)

file_names<-list.files(file.path(root, "raw"))
# fastq file names

mu_site<-matrix(sample(1:genome_size, size=length(file_names)*3), length(file_names),3)
# determine the position on which mutations happen 

test_mutations<-as.data.frame(matrix(NA, length(file_names)*3,10))
colnames(test_mutations)<-c("sample", "position", "ref", "alt", "n_ref", "n_alt", "REF", "ALT", "QUAL", "POS") 
# data frame for writing results

test_mutations[,"sample"]<-rep(file_names, each=3)

system(paste0("mkdir ", root, "/ref_test"))
# make a directory for writing the modified reference genomes

for(i in 1:length(file_names)){

   mu_orig<-rep(NA,3)

   new_fna<-ref_fna
   # duplicate original reference sequence

   for(j in 1:3){  # 1 is bps 2 is deletion 3 is insertion

     mu_orig[j]<-substr(ref_fna, mu_site[i,j],mu_site[i,j])
     # original base

     if(j==1){  # SNPS      
        test_mutations[(i-1)*3+j,"alt"]<-mu_orig[j]
        # original base is then identified as mutated base from the sequencing

        test_mutations[(i-1)*3+j,"ref"]<-sample(setdiff(bases,mu_orig[j]), size=1)
        stri_sub(new_fna, mu_site[i,j],mu_site[i,j])<-test_mutations[(i-1)*3+j,"ref"]
        # replace ref sample base with a new base which will be the reference allele

        test_mutations[(i-1)*3+j,"position"]<-mu_site[i,j]
     }

     if(j==2){ # insert into reference to make DELETIONS

        test_mutations[(i-1)*3+j,"ref"]<-sample(bases, size=1)
        # base to insert

        run<-0
        while(substr(ref_fna, mu_site[i,j]-run,mu_site[i,j]-run)==test_mutations[(i-1)*3+j,"ref"]){run<-run+1}
        # indicate the number of bases prior to the mutation that have the same sequence

        mu_site[i,j]<-mu_site[i,j]-run
        # place the mutation at the start of the run as makes no difference and indexed correctly in vcf

        test_mutations[(i-1)*3+j,"alt"]<-substr(ref_fna, mu_site[i,j], mu_site[i,j])
        # a random base is inserted into the original sequence and so a deletion is identified from the sequencing

        stri_sub(new_fna, mu_site[i,j],mu_site[i,j])<-paste0(test_mutations[(i-1)*3+j,"alt"], test_mutations[(i-1)*3+j,"ref"])
        # add the random base to the ref sample base which will be the reference allele

        test_mutations[(i-1)*3+j,"ref"]<-substr(new_fna, mu_site[i,j],mu_site[i,j]+1)

        test_mutations[(i-1)*3+j,"position"]<-mu_site[i,j]

     }
     if(j==3){ # delete from reference to get INSERTIONS

        run<-0
        while(substr(ref_fna, mu_site[i,j]-run,mu_site[i,j]-run)==mu_orig[j]){run<-run+1}

        # indicate the number of bases prior to the mutation that have the same sequence

        mu_site[i,j]<-mu_site[i,j]-run
        # place the mutation at the start of the run as makes no difference and indexed correctly in vcf

        test_mutations[(i-1)*3+j,"ref"]<-substr(ref_fna,mu_site[i,j],mu_site[i,j])

        test_mutations[(i-1)*3+j,"alt"]<-paste0(test_mutations[(i-1)*3+j,"ref"], mu_orig[j])
        # a random base is deleted from the original sequence and so the original base is identified as an insertion from the sequencing

        if(mu_site[i,2]<mu_site[i,3]){
          adjust<-1  
        }else{
          adjust<-0
        }
        # if a random base was added to the reference before the removal have to increment the position by one

        stri_sub(new_fna, mu_site[i,j]+adjust,mu_site[i,j]+adjust+1)<-test_mutations[(i-1)*3+j,"ref"]
        # delete a random base from ref and so this site will not be present in the ref

        test_mutations[(i-1)*3+j,"position"]<-mu_site[i,j]
     }
  }
  if(mu_site[i,3]<mu_site[i,2]){
    test_mutations[(i-1)*3+2,"position"]<-test_mutations[(i-1)*3+2,"position"]-1
  }
  if(mu_site[i,3]<mu_site[i,1]){
    test_mutations[(i-1)*3+1,"position"]<-test_mutations[(i-1)*3+1,"position"]-1
  }
  if(mu_site[i,2]<mu_site[i,3]){
    test_mutations[(i-1)*3+3,"position"]<-test_mutations[(i-1)*3+3,"position"]+1
  } 
  if(mu_site[i,2]<mu_site[i,1]){
    test_mutations[(i-1)*3+1,"position"]<-test_mutations[(i-1)*3+1,"position"]+1
  } 
   
  fna_path<-file.path(root, "ref_test", paste0(file_names[i], ".fna"))

  cat(header, file=fna_path, sep="\n")
  cat(new_fna, file=fna_path, sep="\n", append=TRUE)
  # write new reference to a file
}
```

#### 5.2 Call simulated mutations

For each library we can map the reads to the new reference genome and call mutations. To achieve this we use the scripts `map_test.sh` and `call_test.sh` which are minor modifications to the scripts `map.sh` and `call.sh`described above. The information is organised and written to test\(\_\)mutations.csv where the columns `ref` and `alt` refer to the simulated mutation and `REF` and `ALT` refer to the mutation called. If a mutation wasn’t called at the correct position `REF` and `ALT` are `NA`.

```
system(file.path(root, "scripts/map_test.sh"))
# map reads to the respective reference

system(file.path(root, "scripts/call_test.sh"))
# call variants with respect to the respective reference

for(i in 1:length(file_names)){

  vcf_path<-file.path(root, "vcf_test", paste0(file_names[i], ".flt.vcf.gz"))
  mini_vcf<-fread(vcf_path, sep = "\t", header = TRUE, skip = "#CHROM")
  # read in vcf

  n_ref<-as.vector(sapply(mini_vcf$INFO, function(x){sum(as.numeric(strsplit(strsplit(x, "DP4=|;MQ=")[[1]][2], ",")[[1]])[1:2])}))
  n_alt<-as.vector(sapply(mini_vcf$INFO, function(x){sum(as.numeric(strsplit(strsplit(x, "DP4=|;MQ=")[[1]][2], ",")[[1]])[3:4])}))

  # number of reads in support of the reference and alternate

  qual<-mini_vcf$QUAL
  # quality score of the call

  pos<-mini_vcf$POS
  # vcf position

  ref<-mini_vcf$REF
  alt<-mini_vcf$ALT
  # called base

  hit<-sapply(test_mutations[(i-1)*3+1:3,"position"], function(x){dist<-abs(x-mini_vcf$POS); if(any(dist<6)){which.min(dist)}else{NA}})
  # find where the known mutations are in the vcf file (if they exist) - match to within 5bp in case there are position shifts
  
  if(any(!is.na(hit))){
    test_mutations[(i-1)*3+1:3,"n_ref"]<-n_ref[hit]
    test_mutations[(i-1)*3+1:3,"n_alt"]<-n_alt[hit]
    test_mutations[(i-1)*3+1:3,"REF"]<-ref[hit]
    test_mutations[(i-1)*3+1:3,"ALT"]<-alt[hit]
    test_mutations[(i-1)*3+1:3,"QUAL"]<-qual[hit]
    test_mutations[(i-1)*3+1:3,"POS"]<-pos[hit]
  }
}

system(paste("rm -r", file.path(root, "vcf_test")))
system(paste("rm -r", file.path(root, "bam_test")))
system(paste("rm -r", file.path(root, "filtered_bam_test")))
system(paste("rm -r", file.path(root, "ref_test")))
```

We can then assess how many mutations are called and whether they pass the strict filter

```
test_mutations$strict<-with(test_mutations, n_alt>=30 & (n_alt/(n_alt+n_ref))>0.95 & QUAL>=20)
# determine those mutations that would pass the strict filter

test_mutations$mtype<-with(test_mutations, c("d","b","i")[nchar(alt)-nchar(ref)+2])

called<-with(test_mutations, table(mtype=mtype, called=!is.na(test_mutations$strict)))
# create a table of called vs. not called across mutation types

called
```

```
##      called
## mtype FALSE TRUE
##     b     0  196
##     d     2  194
##     i     4  192
```

```
strict<-with(test_mutations, table(mtype=mtype, strict=strict))
# create a table of those called mutation passing vs. not the strict filter across mutation types

strict
```

```
##      strict
## mtype FALSE TRUE
##     b     2  194
##     d    38  156
##     i    50  142
```

Almost all mutations are called

Brody, H, A Greener, and CW Hill. 1985. “Excision and Reintegration of the Escherichia Coli k-12 Chromosomal Element E14.” *Journal of Bacteriology* 161 (3): 1112–17.

Danecek, Petr, James K. Bonfield, Jennifer Liddle, John Marshall, Valeriu Ohan, Martin O. Pollard, Andrew Whitwham, et al. 2021. “Twelve years of SAMtools and BCFtools.” *GigaScience* 10 (2): 1–4. https://doi.org/10.1093/gigascience/giab008.

Li, Heng. 2013. “Aligning Sequence Reads, Clone Sequences and Assembly Contigs with BWA-MEM.” *arXiv*, 1303.3997.

Robinson, James T, Helga Thorvaldsdóttir, Wendy Winckler, Mitchell Guttman, Eric S Lander, Gad Getz, and Jill P Mesirov. 2011. “Integrative Genomics Viewer.” *Nature Biotechnology* 29 (1): 24–26.

Zarate, Samantha, Andrew Carroll, Medhat Mahmoud, Olga Krasheninina, Goo Jun, William J. Salerno, Michael C. Schatz, Eric Boerwinkle, Richard A. Gibbs, and Fritz J. Sedlazeck. 2021. “Parliament2: Accurate structural variant calling at scale.” *GigaScience* 9 (12). https://doi.org/10.1093/gigascience/giaa145.
