## Supplementary material for "Fitness effects of new mutations are small and heavily confounded with non-genetic sources of variation in Escherichia coli": Antedependence Model

### Antedependence Models of Environmental Effects

In the discussion we touched on a number of topics that implicitly assumed an antedependence model for the environmental contribution to the between-line variance. Here, we give a short non-technical summary and a more theoretical development of these ideas in order to fix some concepts more precisely.

In what follows we envisage a set-up where a number of MA lines have been propagated for a number of generations from a common ancestor. From each line, a number of sub-lines are then generated and propagated until their fitnesses are measured. These measurements constitute the replicates for each line. We envisage a model for the non-mutational (henceforth environmental) effects whereby new environmental perturbations are constantly being introduced in a (sub)line but their effects decay over time. In what follows two key concepts need to be understood: a) environmental perturbations that arise in a line before the sub-lines are generated will generate between-line variance if they have not decayed by the time measurements are made and b) in the early stages of MA the between-line environmental variance will be small, but as new environmental perturbations are added each generation the environmental variance will increase to some equilibrium value where the input of new environmental perturbations is balanced by the decay of historical perturbations. Concept a) allows us to imagine an experimental design that minimises the environmental between-line variance: if sub-lines can be propagated for as many generations as it takes for their shared environmental effects to decay, any 'memory' of historical environmental perturbations will be lost and there will be no environmental between-line variance, only within-line. In cases where these requirements cannot be met, Concept b) allows us to imagine an alternative experimental design. If the MA lines have been propagated for a sufficiently long period then the environmental variance will have come to an equilibrium and the environmental between-line variance will be constant over subsequent generations. However, the mutational variance will continue to increase each generation (since mutations do not decay) such that *changes* in the between-line variance over the course of an MA experiment can give an estimate of the mutational variance.

To understand these intuitive ideas more formally, we use an antedependence model for the fitness measurement on replicate $j$ for line $i$ taken in generation $t$:

$$y_{ij,t} = g_{i,t-1} + \rho E_{i,t-1} + g^{'}_{i,t} + E^{'}_{i,t} + e_{ij,t} \tag{1}$$

where $g_{i,t-1}$ is the genetic value in the previous generation and $g^{'}_{i,t}$ the increment in the current generation due to new mutations. $E_{i,t-1}$ is the environmental effect in the previous generation that is passed on with fidelity $\rho$ and $E^{'}_{i,t}$ is the increment in this inherited environmental effect. $e_{ij,t}$ is the environmental effect specific to that measurement and includes measurement error. We can rewrite the equation as

$$y_{ij,t} = \sum_{k=0}^{t} g^{'}_{i,k} + \sum_{k=0}^{t} \rho^{t-k} E^{'}_{i,k} + e_{ij,t} \tag{2}$$

The between-line covariance between a set of replicates who had a common ancestor at time $t$ and were measured at time $t+m$ and $t+n$ (i.e the two sub-lines were propagated for $m$ and $n$ generations repsectively) is:

$$Cov(y_{ij,t+m}, y_{ij,t+n}) = \quad tV_m + \rho^{n+m} V_E \sum_{k=1}^{t} \rho^{2(t-k)} \tag{3}$$

assuming that the new mutational and inherited environmental increments are identically and independently distributed, both across lines and in time, with variances $V_m$ and $V_E$ respectively. Here, $g^{'}_{i,0}$ and $E^{'}_{i,0}$ are constant as all lines are assumed to have been derived from the same ancestor at time zero, although this could be relaxed. $tV_m$ is the accumulated mutational variance which grows linearly in time because mutational effects are passed on with perfect fidelity (assuming no directional epistasis). $V_E \sum_{k=1}^{t} \rho^{2(t-k)}$ is the amount of environmental variance that has accumulated by time $t$ and this increases quickly initially, but asymptotes to $V_E/(1-\rho^2)$ as $t$ becomes large. However, the environmental effect initially shared by a set of sub-lines will decay independently in each sub-line as they are propagated, thus reducing their contribution to the between-line covariance by an amount $\rho^{m+n}$. In Figure 1 we give a graphical representation of these equations (see caption for details).

The notation is sufficiently general to cover most MA procedures and measurement protocols. In most large multicelluar organisms $m$ will often be one. For example, an MA line may be propagated by brother-sister mating and at generation $t+1$ a brother and sister are retained as the founders of the next generation and their remaining siblings (the sub-lines) are measured. In most microbial work $m$ may be quite large as a sub-line is grown up and the fitness assays are inherently multigenerational. For example, in our experiment, each sub-line was grown for 16 hours from frozen stock followed by 4 hours on a 96-well plate within which fitness

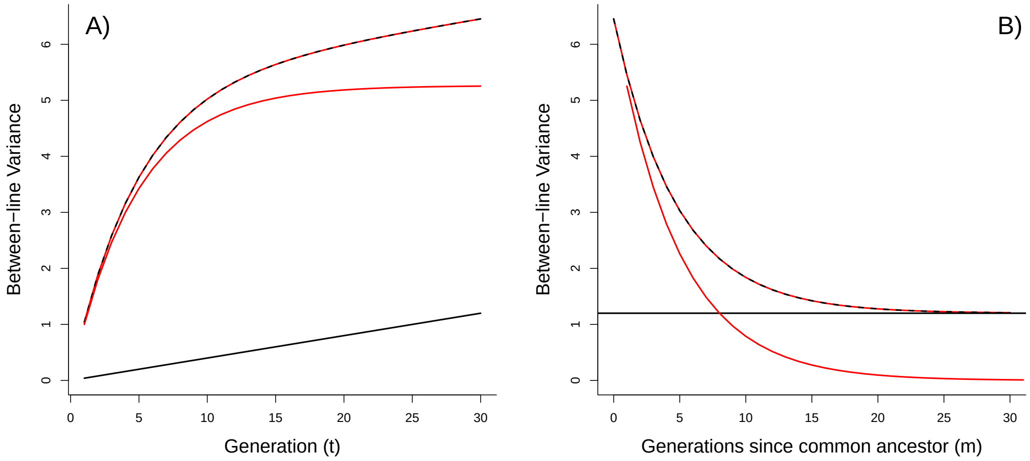


Figure 1: Between-line variances as a function of A) how many generations ($t$) of MA have been conducted assuming no generations separate replicates from their common ancestor ($m = n = 0$) and B) how many generations separate replicates from their common ancestor ($m = n$) given 30 generations of MA ($t = 30$). The dashed black and red line is the total between-line variance which can be decomposed into contributions from the mutational variance (black) and inherited environmental variance (red). The parameters $V_E = 1$, $V_M = 0.04$, $\rho = 0.9$ were used. At $t = 30$ the inherited environmental variance is close to its stationary value $V_E/(1 - \rho^2) = 5.26$.

was measured as the growth rate over a twenty minute period. $m$ is therefore likely to be 10's of generations. At the other extreme, it may be possible to obtain measurements on parents and offspring. Then, $m = 0$ (for the parent) and $n = 1$ (for the offspring) and the between-line (co)variance can be estimated.

If $t$ is sufficiently large, the contribution of the inherited environmental variance to the between-line variance becomes constant, and then the slope of the regression of the between-line variance on generation is a good estimator of $V_m$. However, if the inherited environmental effects have yet to reach stationarity ($t < 15$ in Figure 1) then the slope would overestimate $V_m$ due to the environmental contribution still increasing over time. If $m$ is sufficiently large then the shared environmental effects will have dissipated ($m > 20$ in Figure 1) and no longer contribute to the between-line variance, which is then a good estimator of $V_m$.

If fitness measurements were available at a range of $t$ and/or $m$, yet it was found that neither $t$ or $m$ were sufficiently large for the environmental between-line variance to asymptote (to zero if $m$ is large), the data could still be used to get estimates of $V_m$ by fitting an antedependence model and co-estimating $V_E$ and $\rho$, in addition to $V_M$. Such a model has been developed and fitted to data collected on *Drosophila melanogaster* MA lines (Mackay et al., 1995). However, it should be emphasised that in that paper the non-linear change in the between-line variance is attributed to selection operating on new mutations which essentially causes mutational effects to be inherited without perfect fidelity, as measured through $\rho$. In addition, the authors also note that excess residual heterozygosity in the ancestor (relative to the asymptotic with-line genetic variance expected under MA) and diminishing returns epistasis would also cause the between-line variance to increase faster in early generations, and so interpreting $\rho$ as a measure of selection requires some care. The issue of excess residual heterozygosity, at least, can be dealt with by discarding data from the first $6N_e$ generations of MA, since beyond this the within-line genetic variance will have stabilised at its equilibrium value (where $N_e$ is the effective population size of an MA line).

For completeness, the within-line variance is

$$Var(y_{ij,t+m}|g^{'}_{i,t}, E^{'}_{i,t}) = mV_m + V_E \sum_{k=1}^{m} \rho^{2(m-k)} + V_e \quad (4)$$

where $mV_m$ is the accumulated mutational variance in that replicate, $V_E \sum_{k=1}^{m} \rho^{2(m-k)}$ is the contribution of inherited environmental effects that have arisen in the replicate and $V_e$ the variance in non-inherited environmental effects. Note that if the organism was sexual there would be additional within-line genetic variance due to recombination and segregation.
